# diaPASEF-Powered Chemoproteomics Enables Deep Kinome Interaction Profiling

**DOI:** 10.1101/2024.11.22.624841

**Authors:** Kathryn Woods, Thankhoe A. Rants’o, Alexandria M. Chan, Tanmay Sapre, Grace E. Mastin, Kathleen M. Maguire, Shao-En Ong, Martin Golkowski

## Abstract

Protein-protein interactions (PPIs) underlie most biological functions. Devastating human conditions like cancers, neurological disorders, and infections, hijack PPI networks to initiate disease, and to drive disease progression. Understanding precisely how diseases remodel PPI networks can, therefore, help clarify disease mechanisms and identify therapeutic targets. Protein kinases control most cellular processes through protein phosphorylation. The 518 human kinases, known as the kinome, are frequently dysregulated in disease and highly druggable with ATP-competitive inhibitors. Kinase activity, localization, and substrate recognition are regulated by dynamic PPI networks composed of scaffolding and adapter proteins, other signaling enzymes like small GTPases and E3 ligases, and phospho-substrates. Accordingly, mapping kinase PPI networks can help determine kinome activation states, and, in turn, cellular activation states; this information can be used for studying kinase-mediated cell signaling, and for prioritizing kinases for drug discovery. Previously, we have developed a high-throughput method for kinome PPI mapping based on mass spectrometry (MS)-based chemoproteomics that we named kinobead competition and correlation analysis (kiCCA). Here, we introduce 2^nd^ generation (gen) kiCCA which utilizes data-independent acquisition (dia) with parallel accumulation serial fragmentation (PASEF) MS and a re-designed CCA algorithm with improved selection criteria and the ability to predict multiple kinase interaction partners of the same proteins. Using neuroblastoma cell line models of the noradrenergic-mesenchymal transition (NMT), we demonstrate that 2^nd^ gen kiCCA **(1)** identified 6.1-times more kinase PPIs in native cell extracts compared to our 1^st^ gen approach, **(2)** determined kinase-mediated signaling pathways that underly the neuroblastoma NMT, and **(3)** accurately predicted pharmacological targets for manipulating NMT states. Our 2^nd^ gen kiCCA method is broadly useful for cell signaling research and kinase drug discovery.

## INTRODUCTION

Proteins are the molecular machines that carry out most biological functions. Proteins, however, do not act in isolation, but rather engage in protein-protein interactions (PPIs) to assemble larger complexes that are arrayed into pathways and system-level networks.^1, 2^ Accordingly, the topology of PPI networks can define the functional state of cells and tissues.^3^ Devastating human diseases like cancers, neurological disorders, and infections hijack PPI networks to promote disease initiation and progression, which causes the rewiring of signaling, transcriptional, and metabolic pathways.^4–8^

The 518 protein kinases encoded in the human genome, collectively known as the kinome, control most cellular processes through protein phosphorylation on serine, threonine, and tyrosine (S/T/Y) residues.^9^ Protein phosphorylation can alter a substrate’s activity, cellular localization, stability, and PPIs, which includes kinases themselves. Kinases are frequently dysregulated in human disease, and they are highly druggable with ATP-competitive, small molecule inhibitors.^10–13^ Because of that, kinases have emerged as one of the most important classes of drug targets for combating cancers,^14^ autoimmune and inflammatory diseases,^15, 16^ neurological disorders,^17^ and infectious diseases.^18, 19^ Kinase activity, localization, and substrate recognition are regulated by dynamic PPI networks composed of scaffolding and adapter proteins, other signaling enzymes like small GTPases and E3 ligases, and their phospho-substrates.^20–22^ Profiling kinase PPI networks can, therefore, help determine kinome activation states, and how kinases are connected to signaling pathways.^23, 24^ This information can be used, in turn, for determining the mechanisms of kinase-mediated cell signaling in health and disease, and for prioritizing kinases among the 518 members of the kinome in drug target discovery.^23, 24^ Large-scale maps of PPI networks have been generated using yeast and mammalian two-hybrid systems,^25^ and mass spectrometry (MS)-based approaches like affinity purification (AP)-MS,^26, 27^ size-exclusion chromatography-MS,^28^ protein crosslinking-MS,^29^ and proximity labeling-MS methods like BioID and APEX,^3, 25, 30–33^ however, there is still a critical need for more sensitive and high-throughput methods that can systematically map kinome interaction networks and their dynamics.^23^

We developed the 2^nd^ generation (gen) kinobead competition and correlation analysis (kiCCA) approach for the sensitive and high-throughput profiling of kinome PPIs. Like our 1^st^ gen approach, 2^nd^ gen kiCCA utilizes kinobeads, also known as multiplexed inhibitor beads (MIBs), for affinity purification of the kinome from native cell and tissue lysates, combined with a competitive binding assays using our library of broad-selectivity kinase inhibitors (KIs) that we named kinase interactome probes (KIPs).^23, 34–39^ The competitive binding assay is followed by shotgun nano liquid chromatography (nLC)-MS analysis to determine competed kinases and non-kinase proteins. Correlation analysis then predicts which non-kinase proteins were co-competed with specific kinases, thereby broadly identifying kinase-protein interactions.^23^ To enhance the analytical depth of kinase interactomes, 2^nd^ gen kiCCA relies on data-independent acquisition (dia) – parallel accumulation serial fragmentation (PASEF) for MS data acquisition,^40^ as well as a re-designed CCA algorithm with improved selection criteria and the ability to predict multiple kinase interaction partners of the same proteins. We show **(1)** that kinobead AP-MS with diaPASEF increased the analytical depth of the assayable kinome, which we exploited to determine the kinome-wide selectivity of four clinical anaplastic lymphoma kinase (ALK) inhibitors, and **(2)** that 2^nd^ gen kiCCA increased the number of observable kinome PPIs by 6.1-fold compared to our 1^st^ gen approach, which we utilized to determine signaling pathways and therapeutic targets that promote neuroblastoma (NB) cell phenotypic plasticity.

NB is the most common cancer in infants less than one year of age with 700-800 new cases in the US annually.^41^ NB tumorigenesis is frequently driven by amplification of the transcription factor N-Myc gene (*MYCN*) and activating mutations in the *ALK* gene.^41^ Through improvements in surgery and chemotherapy, low-risk and intermediate-risk NBs are highly treatable with current 5-year survival rates approaching 90%.^41^ High-risk NBs, in contrast, remain a challenge in the clinic and only 50% of patients survive the first 5 years after diagnosis. NB cell phenotypic plasticity can promote therapy resistance and metastasis in high-risk NB, allowing cancer cells to transdifferentiate from a noradrenergic neuronal-like phenotype to a more mesenchymal-like phenotype.^42–46^ For instance, the noradrenergic-mesenchymal transition (NMT) has been shown to increase NB resistance to ALK inhibitor and chemotherapy,^47, 48^ and to modulate the response to disialoganglioside GD2-targeted immunotherapy.^49, 50^ Accordingly, pharmacological strategies to reverse the NMT, and thereby reduce metastasis and therapy resistance, are urgently needed. We used our 2^nd^ gen kiCCA approach to map kinome interaction networks in the noradrenergic neuronal-like NB cell line SH-SY5Y and the isogenic, mesenchymal-like SK-N-SH cell line. This identified specific kinase-mediated signaling pathways that are associated with either of the opposing NMT states. Furthermore, we show that inhibiting these pathways with selective kinase inhibitors can disrupt the opposing NMT phenotypes.

We show that diaPASEF-powered kinobead AP-MS and 2^nd^ gen kiCCA are versatile and effective MS-based chemoproteomic tools for kinome interaction profiling that will be broadly useful for cell signaling research and kinase drug discovery in virtually any disease context.

## RESULTS

### 1. Kinobead AP-MS with diaPASEF enables deep kinome interaction profiling

It has been shown that shotgun nLC-MS analyses of complex peptide mixtures using the diaPASEF method on Bruker timsTOF MS systems improved proteome depth over previous MS systems for high-throughput proteomics.^40^ To determine if such improvements are also observed for relatively low-complexity peptide samples like the ones obtained by affinity purification (AP), we compared the performance of diaPASEF-powered kinobead AP-MS to our previous iterations of the approach.^24, 38, 39^ To determine if diaPASEF improved kinobead AP-MS’s utility for both KI selectivity profiling and kinase-protein interactome profiling, we benchmarked multiple parameters; these included the number of identified and quantified kinases and co-precipitating proteins, and the number of kinases that can be competed off the kinobeads when performing soluble KI binding-competition experiments (**Fig. 1A**).^23, 38, 39^ We subjected the noradrenergic neuronal-like NB cell line SH-SY5Y and the isogenic, mesenchymal-like NB cell line SK-N-SH to kinobead AP-MS analysis, including soluble binding-competition experiments with our 21 KIPs.^23^ Because we have analyzed the two NB cell lines with kinobead AP-MS previously, this allowed us to directly compare performance to our 1^st^ gen approach.^23^ Briefly, peptide samples were analyzed on the Bruker nanoElute 2 – timsTOF Pro 2 nLC-MS system using 45 min nLC gradients and the diaPASEF method with label-free quantification (LFQ, **Fig. 1A**).^40, 51^ We computed raw files using the deep neural network-based search engine Dia-NN 1.8.1 in library-free mode.^52^ This identified and quantified a total 357 kinases in the two NB cell lines, 1.34-fold more compared to the previous iteration of kinobead AP-MS that utilized data-dependent acquisition (dda) on a Thermo Orbitrap Fusion Lumos Tribrid MS system (**Fig. 1B** and **S1A**, and **Table S1**).^23^ We also identified and quantified 6,946 proteins that co-precipitated with the kinobeads in the two NB cell lines, a 2.04-fold increase over our previous workflow (**Fig. 1B**, and **S1A,** and **Table S1**).^23, 38^ This suggested that analyzing relatively low-complexity peptide samples using the diaPASEF method still significantly increased the analytical depth of the kinome and the co-precipitating proteome.

**Figure 1.**
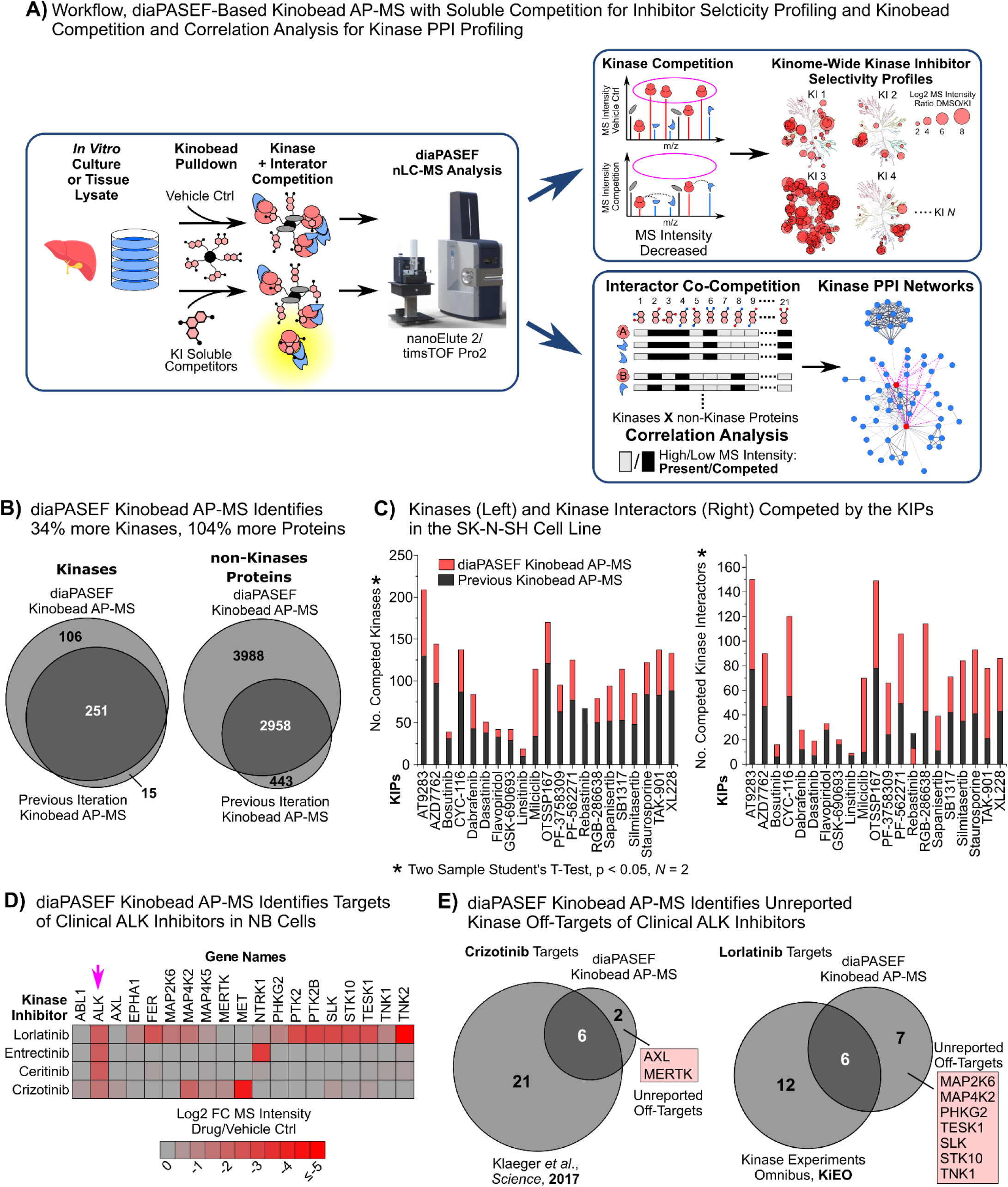
Workflow overview and performance of diaPASEF-powered kinobead AP-MS. **(A)** Detailed overview of our diaPASEF kinobead AP-MS workflow, which enables both kinome-wide KI selectivity profiling and kinase interactome mapping using our kiCCA approach. **(B)** Comparing the number of quantified kinases and non-kinase proteins found in our diaPASEF kinobead AP-MS analysis of the NB cell lines SK-N-SH and SH-SY5Y to our previous kinobead AP-MS approach. **(C)** Number of kinases (left panel) and reported non-kinase interaction partners of these kinases (right panel) that were significantly competed off the kinobeads with our 21 KIPs compared to our previous experiments. Result of diaPASEF kinobead AP-MS assays with soluble competition. Statistics: two-sample Student’s t-test p-value ≤ 0.05, log2 FC ≥ 0, *N* = 2. Refers to Fig. S1B, S1C, and **Table S1**. **(D)** Kinases competed at 1 µM competitor concentration of the ALK inhibitors crizotinib, ceritinib, entrectinib, and lorlatinib as determined by diaPASEF kinobead AP-MS with soluble competition in SK-N-SH cell lysates. Statistics: two-sample Student’s t-test Benjamini-Hochberg (BH)-FDR ≤ 0.05, log2 FC ≥ 0, *N* = 4. Refers to **Table S1**. **(E)** Comparison of significantly competed ALK inhibitor targets from our diaPASEF kinobead AP-MS assay with previously reported targets of these inhibitors. Statistics: see **(D)**. Refers to **Table S1**.

Next, we compared the number of kinases that could be competed with our 21 KIPs to our previous kinobead AP-MS experiments.^23^ This showed an average 1.43-fold increase in competed kinases across all KIPs (**Fig. 1C, S1B, S1C**, and **Table S1**). We also determined if diaPASEF identified more previously reported interaction partners of kinases that were co-competed with the 21 KIPs.^23, 53^ This showed that we identified 447 reported kinase interaction partners in the two NB cell lines that were co-competed with the kinase targets of the respective KIP, an average 1.86-fold increase across all KIPs (**Fig. 1C** and **S1C,** and **Table S1**).^23^ Collectively, these results showed that diaPASEF-powered kinobead AP-MS with soluble competition significantly expanded the number kinases that are assayable in KI binding-competition experiments and identified a larger number of kinase interaction partners for deep kinome PPI profiling.

To clarify if the increased analytical depth of diaPASEF-powered kinobead AP-MS can identify unreported kinase inhibitor targets, we profiled the kinome selectivity of the 1^st^ gen ALK inhibitor crizotinib, the 2^nd^ gen ALK inhibitor ceritinib, and the 3^rd^ gen ALK inhibitors entrectinib and lorlatinib.^54^ Several ALK inhibitors are approved for the treatment of non-small cell lung cancers (NSCLCs) that carry *ALK* gene fusions.^55^ Furthermore, the *ALK* gene is affected by gain of function mutations in 14% of high-risk NB cases, and ALK inhibitors like lorlatinib are being tested in clinical trials against high-risk NB.^56^ A better understanding of ALK inhibitor target profiles in NB cells will help determine their mechanisms of action, optimize treatments, and avoid off-target toxicities.

Accordingly, we profiled the kinome-wide selectivity of crizotinib, ceritinib, entrectinib, and lorlatinib in SK-N-SH cell lysate at 1 µM KI competitor concentration. This showed that entrectinib (3^rd^ gen) exclusively bound to its main targets ALK and the high-affinity nerve growth factor receptor (NTRK1),^57^ and that ALK was the sole interactor of ceritinib (2^nd^ gen),^58^ thus confirming their high selectivity across the kinome (**Fig. 1D**, **Table S1**). In contrast, crizotinib (1^st^ gen) and lorlatinib (3^rd^ gen) showed binding to several off-target kinases that have not been reported previously (*KI*nase Experiments Omnibus, KiEO, http://kieo.tanlab.org).^36^ Thus, we found that crizotinib also bound to the receptor tyrosine kinases AXL and Mer (MERTK) that can promote cancer cell survival and therapy resistance.^59, 60^ In addition to known off-targets like the tyrosine kinases TNK2, PTK2, and PTK2B (http://kieo.tanlab.org),^61^ lorlatinib showed strong off-target binding to several serine-threonine kinases, including MAP4K2 and MAP2K6, and the non-receptor tyrosine kinase TNK1, which has not been reported previously. TNK1 has recently been shown to promote tumor growth and autophagy, and MAP4K2 and MAP2K6 activate the JNK and p38 branches of mitogen-activated protein kinase (MAPK) signaling that can confer therapy resistance to cancer cells.^62, 63^ Collectively, our results identified previously unreported off-targets of clinical ALK inhibitors, whose inhibition may contribute to their clinical efficacy in NSCLCs and NBs.

### 2. Improved selection criteria for CCA boost the depth of kinome PPI analyses

Previously, we observed that inconsistently and weakly competed kinases in our kinobead AP-MS binding-competition experiments produced false-positive PPI predictions in our CCA, and that we had to remove such kinases from our CCA analysis to prevent the accumulation of false positives.^23^ To more systematically remove kinases that may produce false positives from our current analyses, we performed parameter scanning, selecting kinases that fulfill increasingly stringent t-test p-value and log2 MS intensity fold-change (FC) cut-off criteria when performing binding-competition experiments with our 21 KIPs (**Fig. 2A** and **S2A**). To determine if inconsistently or weakly competed non-kinase proteins lead to the accumulation of false-positive PPI predictions as well, we performed the same parameter scanning experiments for non-kinase proteins (**Fig. 2A** and **S2A**). To estimate the performance of our CCA in predicting the largest possible number of true vs. false PPIs when using specific cut-off criteria for input kinases and non-kinase proteins, we introduced a CCA score (S_CCA_) that incorporated the difference in the median of CCA Pearson’s r-values for reported vs. unreported kinase PPIs in the BioGRID database, multiplied with the number of reported kinase PPIs at each cut-off (S_CCA_ = (median r reported – median r unreported) * *N*_reported_).^53^ This showed that an increasingly stringent log2 MS intensity FC cut-off for kinases led the S_CCA_ to undergo a maximum at ∼2.5-fold for both the SK-N-SH and the SH-SY5Y cell line (**Fig. 2A** and **S2A**). In contrast, an increasingly stringent FC cut-off for non-kinase proteins did not cause the S_CCA_ to undergo another maximum (**Fig. 2A** and **S2A**). The same was true for increasingly stringent *p*-value cut-offs (**Fig. 2A** and **S2A)**. We concluded that more stringent selection criteria for input kinases, but not non-kinase proteins, ensured that our CCA predicted most true kinase PPIs correctly, while keeping the number of false-positive kinase PPI predictions at a minimum. To identify an optimal cut-off for kinases to input into CCA that considers both the log2 MS intensity FC and the t-test *p*-value for KIP competition, we next plotted the S_CCA_ for different FC and p-value cut-offs against one another (**Fig. 2B** and **S2B)**. This showed that the S_CCA_ rapidly plateaued at a log2 FC ≥ 0.5 and a p-value of ≤ 0.05 for CCA analysis of the SK-N-SH cell line, and at a log2 FC ≥ 0.5 and a p-value of ≤0.01 for CCA analysis of the SH-SY5Y cell line. To clarify if these new selection criteria improved performance over our 1^st^ gen CCA workflow, we applied our previous cut-off criteria for both kinases and non-kinases (log2 MS intensity FC >0.75 and t-test p-value < 0.05) to our diaPASEF kinobead AP-MS data from the two NB cell lines and performed CCA. We then compared the number and identity of reported kinase PPIs that were predicted by our 2^nd^ gen CCA to the number of PPIs that were predicted by our 1^st^ gen CCA.^23^ This showed that using our newly defined selection criteria led the CCA to correctly predict 81% and 47% more previously reported kinase PPIs in the SK-N-SH and SH-SY5Y cell lines, respectively (**Fig. 2C**).

**Figure 2.**
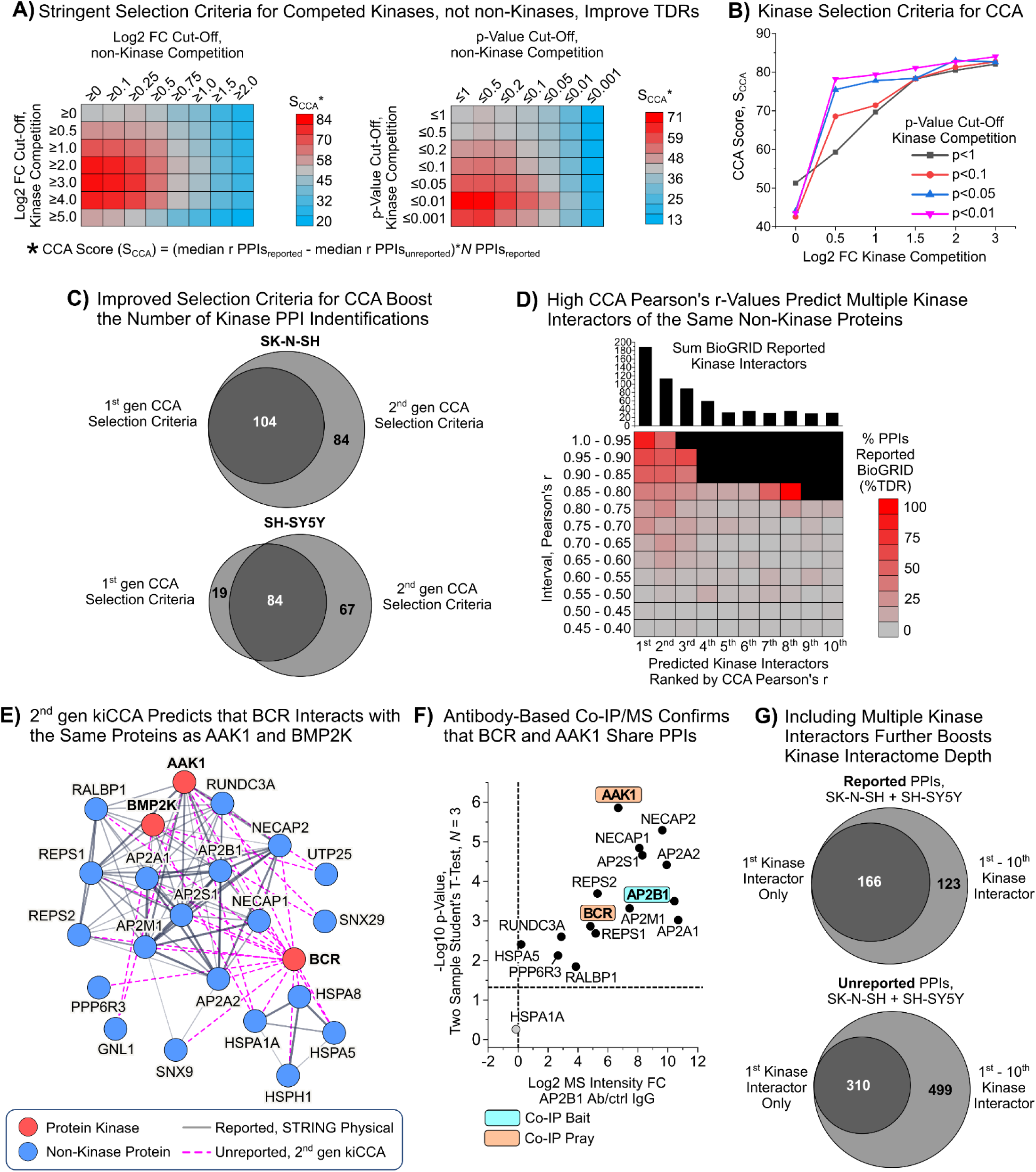
Re-defining the selection criteria for the CCA. **(A)** Parameter scanning experiment determining the CCA score (SCCA) for kinase PPIs predicted by our CCA in the SK-N-SH cell line. Increasingly stringent Student’s t-test log2 FC and p-value cut-off criteria were applied and only kinases or non-kinases fulfilling these criteria were used for CCA. Statistics: two sample Student’s t-test, range of log2 FC ≥ 0 to ≤ 5, and p ≥ 0 to ≤ 0.001. Refers to **Fig. S2A**. **(B)** Plotting the SCCA for varying log2 FC cut-offs and p-Value cut-offs against one another identified optimal mixed cut-off criteria for kinases input into our CCA. Refers to **Fig. S2B**. **(C)** Venn diagrams showing the overlap of reported kinase PPIs that were predicted by our 1^st^ gen CCA algorithm and PPIs predicted by our 2^nd^ gen CCA algorithm. Both 1^st^ and 2^nd^ gen CCA used the same diaPASEF kinobead AP-MS data from the two NB cell lines SK-N-SH and SH-SY5Y as the input. Refers to **Table S1**. **(D)** Heatmap showing the %TDR for PPIs achieved by our 2^nd^ gen CCA separated into 0.05-unit intervals of the CCA Pearson’s r-value for the 1^st^ through 10^th^ predicted kinase interactor of a non-kinase protein. The %TDR reflects the percentage of BioGRID-reported divided by BioGRID-unreported PPIs. Refers to **Fig. S2C**. **(E)** STRING network for interaction partners of the kinases AAK1, BMP2K, and BCR, as predicted by 2^nd^ gen CCA. **(F)** Volcano plot showing the results of a co-IP/MS experiment using a specific antibody that binds AP2B1. Only proteins that are also members of the STRING interaction network in (E) are shown. Statistics: two sample Student’s t-test p < 0.05. Refers to **Fig. S2E**. **(G)** Venn diagrams comparing the total number of predicted kinase PPIs of 1^st^ predicted kinase vs. 1^st^ – 10^th^ predicted kinase, as was determined by 2^nd^ gen CCA in the NB cell lines SK-N-SH and SH-SY5Y.

### 3. Second generation kiCCA identifies multiple kinase interactors of the same proteins

Signaling proteins, enzymes, and transcription factors often interact with, and are phosphorylated, by multiple kinases, enabling signal integration and introducing redundancy to upstream kinase pathways.^64, 65^ Determining kinase pathway redundancy, in turn, is critical for understanding cell signaling mechanisms and to predict responses to KI therapy.^35, 66^ The previous iteration of our kiCCA workflow was able to predict only the kinase-protein interactor pair with the highest CCA Pearson’s r value, and thus was unable to identify multiple kinase interactors of the same protein. To address this shortcoming, we re-computed our 2^nd^ gen kiCCA data, determining kinases with the 1^st^ to 10^th^ highest CCA Pearson’s r-value for all quantified non-kinase proteins.

We then mapped CCA PPI predictions against reported kinase PPIs in the BioGRID interaction database, calculated %TDRs for interactions with the 1^st^ to 10^th^ kinase (%TDR = (*N*_reported_/*N*_unreported_) * 100), and plotted %TDRs for 0.05 Pearson’s r intervals on a heat map (**Fig. 2D** and **S2C**).^53^ These %TDR maps allowed us to convert CCA Pearson’s r-values into TDR estimates for the 1^st^ to 10^th^ predicted kinase interactors of non-kinase proteins (**Table S1**). For the 1^st^ kinase interactor, %TDRs approached 100% for Pearson’s r-values >0.95 and monotonously decreased to ∼10% for Pearson’s r-values between 0.65 and 0.6. We observed the same trend for the 2^nd^ and 3^rd^ kinase interactor, and in some cases observed >10% TDRs for up to the 10^th^ kinase interactor for some broad specificity signaling adapters like 14-3-3 family proteins. For instance, 2^nd^ gen kiCCA predicted 7 reported and 9 unreported kinase interactors of the 14-3-3 proteins YWHAB, YWHAE, YWHAG, YWHAH, YWHAQ, and YWHAZ (**Fig. S2D** and **Table S1**). Likewise, 2^nd^ gen kiCCA predicted 4 reported and 6 unreported kinase interactors of the signaling adapter growth factor receptor-bound protein 4 (GRB4 or NCK2) that broadly binds trysosine-phosporylated receptor and non-receptor tyrosine kinases (**Fig. S2D** and **Table S1**). These results suggested that 2^nd^ gen kiCCA can correctly predict multiple kinase interaction partners of the same proteins.

Among the multiple kinase interactors predicted to interact with the same proteins (%TDR > 10), we found that many are highly homologous kinases like the hippo kinases STK3 and STK4, and casein kinase catalytic subunits α1 and α2 (CSNK2A1 and A2, **Table S1**). This reflected both the structural similarity of these kinases, and their overlapping biological functions. We also identified structurally dissimilar kinases that were predicted to interact with the same proteins. For instance, 2^nd^ gen CCA predicted that the kinase breakpoint cluster region protein (BCR) interacted with the same proteins as the adapter-associated kinase 1 (AAK1) and the BMP-2-inducible protein kinase (BMP2K) in SK-N-SH cells (**Fig. 2E** and **Table S1**). To validate that BCR interacted with same proteins as AAK1 and BMP2K, we performed co-IP/MS experiments using a selective antibody targeting the common predicted interactor adapter protein 2 (AP2) subunit β1 (AP2B1) in SK-N-SH cell lysate. This revealed that AP2B1 co-precipitated both AAK1 and BCR, and 12 additional members of the network, but not BMP2K (**Fig. 2F** and **Table S2**). This suggested that AAK1 and BCR, but not BMP2K, are part of the same PPI network. To clarify if AAK1 and BCR are part of the same protein complex, we performed another co-IP/MS experiment using a selective antibody targeting AAK1. This showed that AAK1 co-precipitated BCR and 7 additional members of this network (**Fig. S2E** and **Table S2**). These results validated that 2^nd^ gen kiCCA can correctly predict multiple kinase interaction partners of the same proteins.

Comparing the total number of predicted kinase PPIs at a %TDR > 10 when only considering the 1^st^ kinase interactor vs. including the 1^st^ through the 10^th^ interactor, we observed 74% more reported and 261% more unreported PPIs across the two NB cell lines for a total of 1,098 unique kinase-protein interactions (**Fig. 2G** and **Table S1**); this presents an 6.1-fold increase in unique kinase PPIs compared to our 1^st^ gen kiCCA of the two NB cell lines (*N* = 179 predicted PPIs). This showed that including multiple kinase interactors greatly expanded observable kinome interactions.

### 4. A detailed map of NMT-dependent kinome PPI rewiring in neuroblastoma

Having established a map of the kinome and its PPIs in the isogenic mesenchymal-like SK-N-SH cell line and the noradrenergic neuronal-like SH-SY5Y cell line (**Fig. 3A**), we next asked how the NB cell NMT alters the abundance of the kinome and its PPIs. Kinases that interact with signaling complexes in an NMT-dependent manner may promote phenotypic switching and present druggable targets to inhibit NB phenotypic plasticity, metastasis, and therapy resistance.^48–50^ Accordingly, we performed differential expression analysis (DEA) of kinases and their interaction partners between the two NB cell lines as described previously.^23^ This identified 302 kinases and 358 kinase interaction partners that were differentially abundant between the two NB cell lines (**Fig. 3B** and **Table S2**), an increase of 58% and 295% compared to our previous experiments.^23^ Concordant with our previous results,^23^ kinases that are typically expressed in the central and peripheral nervous system like ALK and the peripheral plasma membrane protein CASK were highly abundant in the noradrenergic neuronal-like SH-SY5Y cell line (**Fig. 3B**). In contrast, kinases typically expressed in the mesenchymal lineage like the receptor tyrosine kinases AXL and MET, and the TGFβ receptor type-2 (TGFBR2) were highly enriched in the SK-N-SH cell line. This validated that the two NB cell lines exist in opposing NMT states. Notably, our 2^nd^ gen approach identified 133 kinases that showed differential abundance between the NB cell lines that we did not detect using our 1^st^ gen approach. These included kinases controlling neuronal growth, differentiation and survival like the neurotrophic tyrosine kinase receptors type 1 and 2 (NTRK1 and 2), cell polarity like the MAP/microtubule affinity-regulating kinases 1-4 (MARK1-4), and cell migration like the myosin light chain kinase (MYLK, **Fig. 3B** and **Table S2**), thus yielding additional information on how these processes may be controlled by kinases during the NMT.

**Figure 3.**
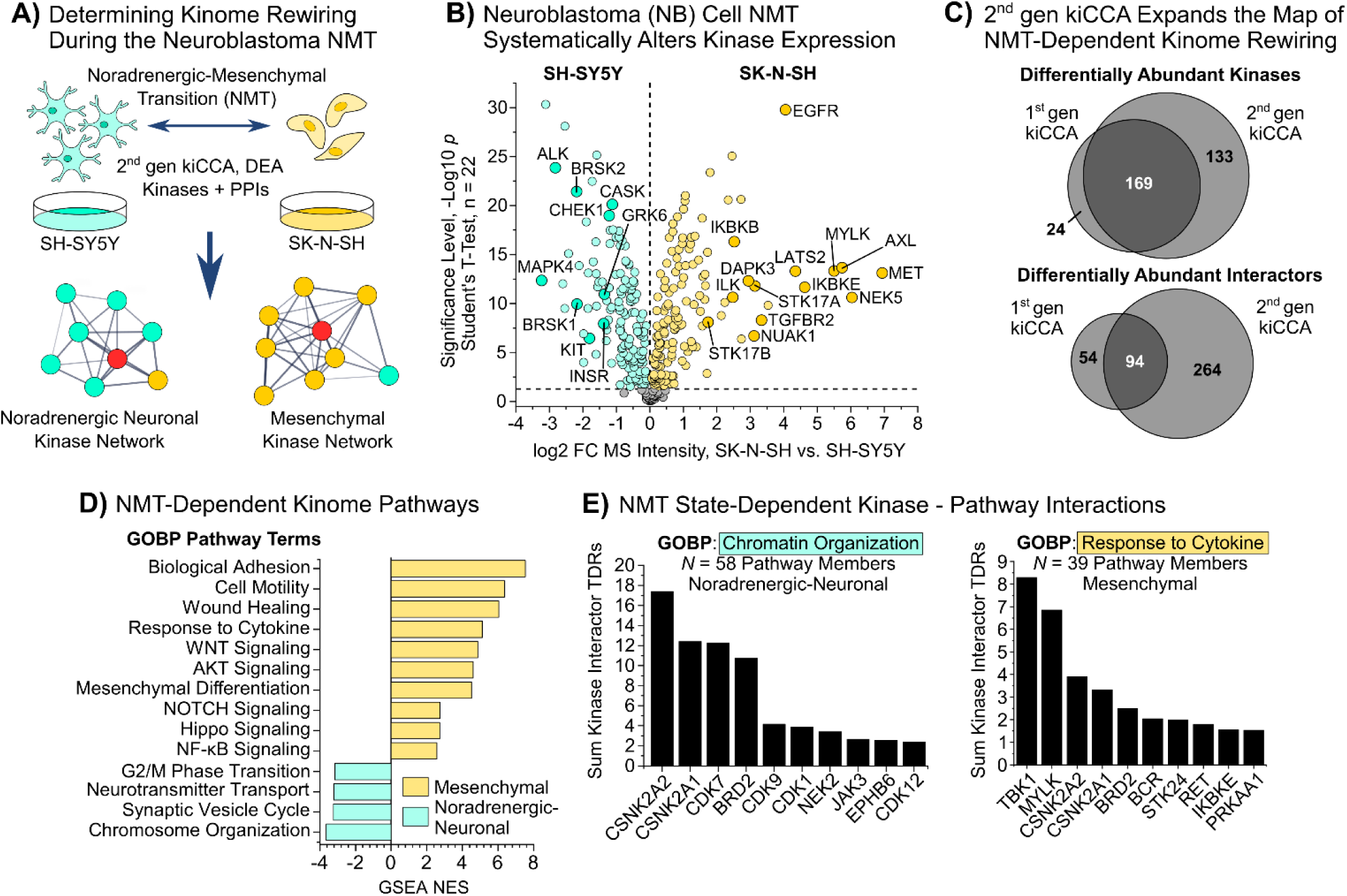
Mapping changes in kinome PPIs that are associated with the NB cell NMT using our 2^nd^ gen kiCCA approach. **(A)** Schemata: the noradrenergic-neuronal SH-SY5Y cell line and the isogenic mesenchymal-like SK-N-SH cell line serve as *in vitro* model for the NB cell NMT. **(B)** Volcano plot showing the differential abundance of kinases between the two opposing NMT states represented by the SH-SY5Y and SK-N-SH cell line. Statistics: one sample Student’s t-test, BH-FDR < 0.05, *N* = 22. Refers to **Table S2**. **(C)** Venn diagrams comparing the differential abundance of kinases and kinase interaction partners between the two NMT states as determined by either 1^st^ gen or 2^nd^ gen kiCCA (for statistics, see (B)). Refers to **Table S2**. **(D)** Results of a GSEA with GOBP terms applied to the results of differential expression analysis of kinase and kinase interaction partners between the two NB cell line SH-SY5Y and SK-N-SH; only pathways that achieved an FDR < 0.05 are shown (for statistics, see (B)). **(E)** Association of kinases with pathways through the sum of %TDRs of their interaction partners that have been associated with specific GOBP pathways terms shown in panel (D). NES is the normalized enrichment score. Refers to Fig. 3A and **Table S1**.

The 358 differentially abundant kinase interactors were part of 168 kinase PPI networks that control diverse cellular processes (2^nd^ gen kiCCA interaction %TDR > 10, **Table S2**). Applying gene set enrichment analysis (GSEA) to the DEA results of kinases and their interaction partners confirmed that pathways related to the mesenchymal phenotype were upregulated in the SK-N-SH cell line. These included pathways related to cell motility, wound healing, cytokine signaling, including WNT and NF-κB signaling, and NOTCH and Hippo signaling (**Fig. 3D** and **Table S2**).

In contrast, pathways enriched in the noradrenergic neuronal-like SH-SY5Y cell line included terms related to central nervous system function like learning, synaptic vesicle cycle, and neurotransmitter transport, as well as terms related to chromatin regulation and cell cycle progression (**Fig. 3D** and **Table S2**). To determine which kinases interacted with the pathways that were associated with NMT phenotypes, we systematically mapped non-kinase pathway members to specific kinases for 14 representative pathways terms (**Fig. 3E** and **S3A**, and **Table S2**). This created a map of kinase-pathway interactions in the NMT, suggesting both reported and unreported kinase functions in phenotypic plasticity. For instance, as we described previously, casein kinase 2 (CK2) interacted with chromatin remodeling complexes in the noradrenergic neuronal-like SH-SY5Y cell line. In addition, we found here that TANK-binding kinase 1 (TBK1), AAK1, and STE20-like kinase MST3 (STK24) interacted preferentially with proteins that control NF-κB, WNT, and NOTCH signaling in the mesenchymal-like SK-N-SH cell line (**Fig. 3E** and **S3A**, and **Table S2**). This suggested that inhibiting these kinases may reprogram the opposing NMT states.

### 5. CK2 and TBK1 present target candidates for reprogramming NB cell phenotypic states

NB cell phenotypic transitions like the NMT can promote metastasis, disease relapse, and therapy resistance.^42–44^ We sought to determine if our 2^nd^ gen kiCCA approach can prioritize kinase targets for manipulating NMT phenotypic states. We focused on kinase complexes that showed NMT state-dependent changes in their composition, particularly kinase complexes that control chromatin organization and cytokine signaling; this is because chromatin remodeling plays critical roles in cancer cell phenotypic plasticity.^67^ Likewise, cell signaling pathways that are triggered by cytokines like TGFβ, WNT5α, interleukins, and TNFα have been shown to promote cancer cells’ transitions to a mesenchymal-like state.^68^

Concordant with our previous results, we found that CK2 interacted with members of the polycomb repressive complex 1 (PRC1), including the autism susceptibility gene 2 protein (AUTS2), preferentially in the SH-SY5Y cell line (**Fig. 4A** and **Table S2**).^23^ AUTS2 has been shown previously to activate the transcription of neuronal genes in cooperation with CK2, suggesting that PRC1 complexes that contain CK2 and AUTS2 can promote the noradrenergic neuronal-like phenotype in NB cells.^69^ A function for CK2 and AUTS2 in the NB cell NMT has not been reported previously, however. Supporting our hypothesis, 2^nd^ gen kiCCA identified additional PRC1 complex members that interacted with CK2 preferentially in the SH-SY5Y cell line, and that can activate the expression of neuronal genes (**Fig. 4A** and **Table S2**); these included fibrosin-1-like protein (FBRSL1, also known as AUTS2L), the RING1 and YY1-binding protein (RYBP), and the chromobox protein homolog 2 (CBX2).^69, 70^ These results led us to hypothesize that pharmacological inhibition of CK2 can destabilize the noradrenergic neuronal-like phenotype in the SH-SY5Y cell line.

**Figure 4.**
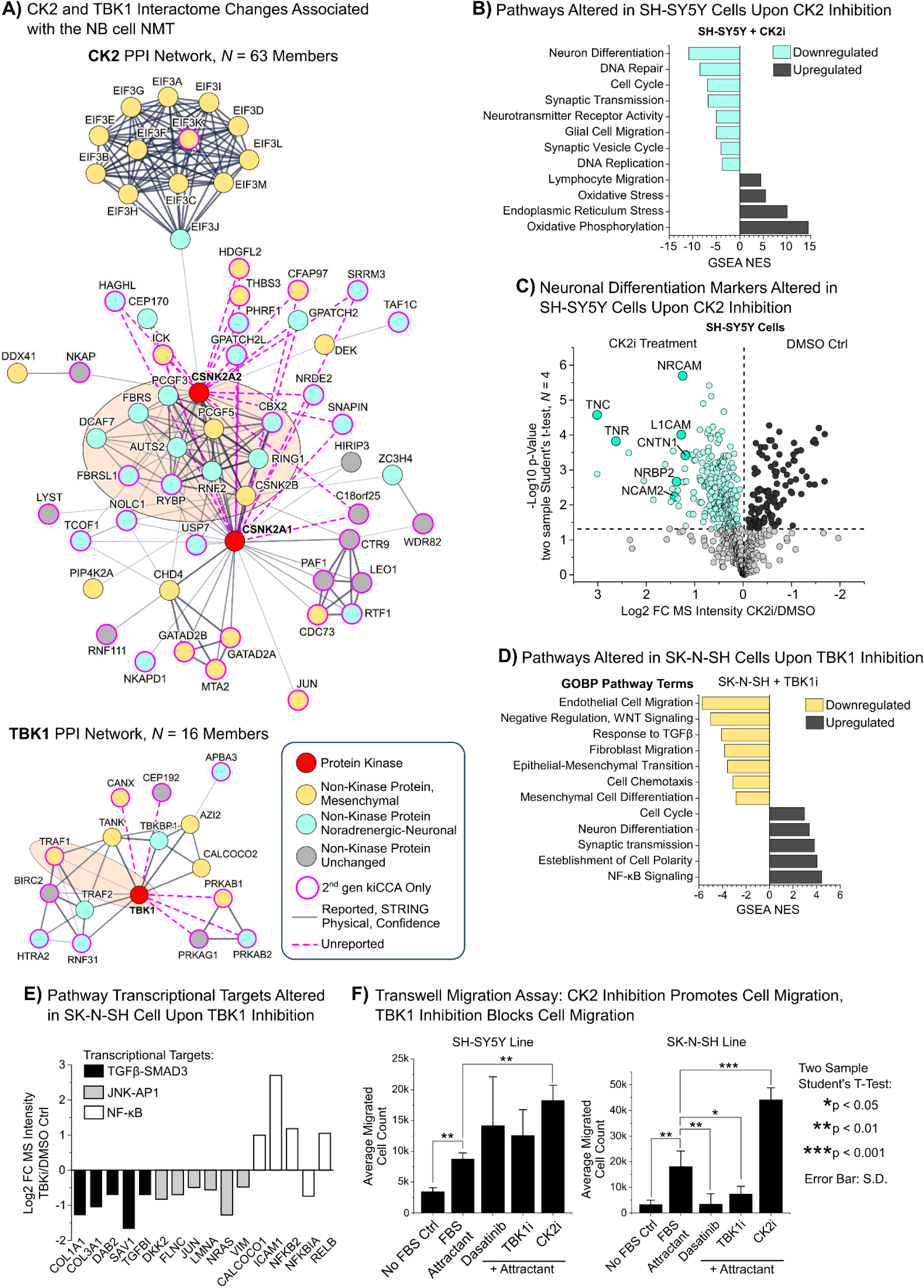
Utilizing 2^nd^ gen kiCCA to prioritize kinase targets for pharmacologically manipulating NB cell NMT states. **(A)** STRING interaction networks (v12.0) of kinase PPIs predicted by 2^nd^ gen kiCCA in the isogenic NB cell lines SK-N-SH (mesenchymal-like) and SH-SY5Y (noradrenergic neuronal-like). Only physical interactions were used to build the STRING network. Relates to **Table S1**. **(B)** Results from GSEA of global protein expression data, comparing SH-SY5Y cells treated with either the CK2 inhibitor SGC-CK2-1 or DMSO (vehicle) for 4 days. Only pathway terms that achieved an FDR < 0.05 are shown. Relates to **Table S2**. **(C)** Volcano plot showing expression differences of neuronal marker proteins between SH-SY5Y cells treated with either the CK2 inhibitor SGC-CK2-1 or dimethyl sulfoxide (DMSO, vehicle) for 4 days. Proteins included in the GOBP gene set ‘Neuronal Differentiation’ were used to define proteins as neuronal marker proteins. Statistics: two sample Student’s t-test, BH-FDR < 0.05, *N* = 3 Relates to **Table S2**. **(D)** Results from GSEA of global protein expression data, comparing SK-N-SH cells treated with either the TBK1 inhibitor GSK8612 or DMSO (vehicle) for 4 days. Only pathway terms that showed an FDR < 0.05 are shown. Relates to **Table S2**. **(E)** Differences in the expression of proteins that are the transcriptional targets of TGFβ-SMAD3, JNK-AP1, and NF-κB signaling between SK-N-SH cells treated with either the TBK1 inhibitor GSK8612 or DMSO (vehicle) for 4 days. Result of global proteome profiling. All proteins shown significantly differed in expression. Statistics: two sample Student’s t-test, BH-FDR < 0.05, *N* = 4. Refers to **Table S2**. **(F)** Trans well migration assay showing that CK2 inhibition with SGC-CK2-1 promoted migration in both SH-SY5Y cells and SK-N-SH cells, and that TBK1 and tyrosine kinase inhibition using GSK8612 and dasatinib, respectively, significantly inhibited cell migration only in the SK-N-SH cell line. Statistics: two sample Student’s t-test p < 0.05, *N* = 4; error bars are the S.D.

2^nd^ gen kiCCA also revealed that TBK1 was the most prominent kinase interacting with proteins that control cytokine signaling. Particularly, the interaction of TBK1 with the TNF receptor associated factor 1 (TRAF1) was highly abundant in the mesenchymal-like SK-N-SH cell line (**Fig. 4A** and **Table S2**). TBK1 controls interferon and NF-κB signaling in a context-dependent manner.^71^ TRAF1 is a scaffolding protein that can activate NF-κB and JNK signaling, which have both been shown to promote the mesenchymal differentiation of cancer cells.^72–75^ Furthermore, TRAF1 can recruit the anti-apoptotic E3 ligases BIRC2 and BIRC3 to TNF receptors, two proteins that we also found to be part of the TBK1 PPI network (**Fig. 4A**).^76^ This suggested that the TBK1- TRAF1 complex promoted the mesenchymal-like state in the SK-N-SH cell line, and that pharmacological inhibition of TBK1 can destabilize the mesenchymal-like phenotype.

To test our target hypotheses, we treated the SH-SY5Y (noradrenergic neuronal-like) and SK-N-SH (mesenchymal-like) cell lines with the selective CK2 inhibitor SGC-CK2-1 and the selective TBK1 inhibitor GSK8612 for four days.^77, 78^ To determine inhibitor effects on NMT pathways, we then performed global MS-based proteome analysis; this showed that CK2 inhibition significantly altered the expression of 3,202 proteins in the SH-SY5Y cell line (40% of the proteome) and the expression of 2,147 proteins in the SK-N-SH cell line (26% of the proteome), whereas TBK1 inhibition did not alter protein expression in the SH-SY5Y cell line, but significantly altered the expression of 708 proteins in the SK-N-SH cell line (8.5%, **Table S2**, two sample student’s t-test, BH-FDR ≤ 0.05, *N* = 4 and *N* = 3). This suggested that CK2 broadly controls gene expression or protein stability in both the SH-SY5Y and SK-N-SH cell lines, and that TBK1 does so only in the SK-N-SH cell line. Applying GSEA with GOBP pathway terms to the global proteome data showed that CK2 inhibition downregulated pathways related to neuronal function, including neuron differentiation and synaptic transmission, as well as DNA repair and cell cycle terms, and upregulated pathways related to cell migration, oxidative phosphorylation, and stress in both NB cell lines (**Fig. 4B, Fig. S3B,** and **Table S2**). Downregulation of neuronal pathways was evident also at the level of individual neuronal markers. Thus, we observed the systematic downregulation of, e.g., the neuronal adhesion molecules NRCAM, NRCAM2, and L1CAM, as well as the neural extracellular matrix proteins tenascin-C (TNC) and tenascin-R (TNR, **Fig. 4C**).

In contrast, in SK-N-SH cells, TBK1 inhibition downregulated pathways related to cell migration, TGFβ signaling, pathways that negatively regulate canonical WNT signaling, and mesenchymal differentiation (**Fig. 4D** and **Table S2**). Canonical NF-κB signaling, on the other hand, was upregulated, hinting that TBK1 inhibits this pathway in mesenchymal NB cells. Analyzing the expression of individual pathway marker proteins confirmed that TGFβ signaling became downregulated and NF-kB signaling upregulated in response to TBK1 inhibition (**Fig. 4E**). Concordant with TRAF1 functions in c-Jun N-terminal kinase (JNK) signaling, we also observed that the transcription factor JUN itself and several AP1 (JUN-FOS) targets became downregulated (**Fig. 4E**); this included the mesenchymal markers vimentin (VIM) and prelamin-A/C (LMNA), as well as the NB oncogenic driver GTPase NRas (NRAS). We concluded that that the TBK1-TRAF1 complex may balance NF- κB inhibition with TGFβ and JNK-AP1 pathway activation to promote the mesenchymal state in NB cells.

Next, to test if changes in molecular pathways correlate with changes in cellular phenotypes, we performed trans-well assays, determining changes in cell migration, a hallmark of the transition to a more mesenchymal-like state. Our GSEA showed that inhibiting CK2 upregulated pathways related to cell migration, whereas TBK1 inhibition downregulated cell migration pathways. Indeed, we observed that CK2 inhibition significantly increased cell migration both in the SH-SY5Y and the SK-N-SH cell line (**Fig. 4F**), suggesting that both cell lines became relatively more mesenchymal-like. In contrast TBK1 inhibition significantly decreased migration in SK-N-SH cell lines, as did dasatinib, a broad selectivity inhibitor of Src-family kinases which served as our positive control (**Fig. 4F**). This confirmed that TBK1 inhibition antagonizes mesenchymal-like traits.

Collectively, our results demonstrated that 2^nd^ gen kiCCA can be used to prioritize kinase targets for altering NB cell NMT states, which in turn could be exploited to modulate NB responses to targeted therapy and to inhibit metastasis and disease relapse.

## DISCUSSION

Here, we demonstrated that diaPASEF-powered kinobead AP-MS and 2^nd^ gen kiCCA are versatile and efficient MS-based chemoproteomic tools for kinome interaction profiling, which includes kinase-inhibitor and kinase-protein interactions. Specifically, we showed that diaPASEF kinobead AP-MS identified previously unreported kinase targets of clinical ALK inhibitors that may explain their mechanism of action and clinical efficacy. Thus, lorlatinib may inhibit kinases that are involved in JNK and p38 signaling, and in autophagy (TNK1), all processes that have been shown to promote therapy resistance *in vivo*.^62, 63, 79^ Lorlatinib is the ALK inhibitor that has progressed the furthest in clinical trials against high-risk NB to date,^56^ and has the most off-target kinases of all ALK inhibitors that we tested. Collectively, this suggested that the polypharmacology of lorlatinib is critical for its clinical efficacy, and that more efficient inhibitors with engineered polypharmacology could be developed utilizing the insights that we obtained.

We also showed that our 2^nd^ gen kiCCA approach identified 1,098 unique kinase PPIs in the two isogenic NB cell lines SK-N-SH and SH-SY5Y, 74% of which have not been reported previously.^53^ Comparing kinase interactome changes between the mesenchymal-like SK-N-SH cell line and the noradrenergic neuronal-like SH-SY5Y revealed 168 kinases that showed differences in their PPIs between the opposing NMT states. We utilized this information for target prioritization, and we correctly predicted kinases that when inhibited, destabilize the respective NMT states. Thus, concordant with our target hypotheses, inhibiting CK2 destabilized the noradrenergic neuronal-like phenotype, while inhibiting TBK1 destabilized the mesenchymal phenotype. Intuitively, this suggested that TBK1 inhibition may be useful to sensitize NBs to therapy and to prevent metastasis. Yet, inhibiting CK2 to destabilize the noradrenergic neuronal-like phenotype may be of translational value as well. Thus, recent studies suggested that noradrenergic neuronal-like NB cells are more resistant to GD2-targeted immunotherapy than their mesenchymal-like counterparts.^50^

Collectively, our results present compelling evidence that mapping alterations of kinase PPI networks in disease can be used to prioritize targets for therapeutics development and provide a deep knowledgebase of kinase signaling pathways that are involved in the NB cell NMT. This knowledgebase contains several additional leads on kinases that may control NB cell NMT states; these include AAK1, specific Src-family kinases like CSK, and the myosin light chain kinase (MYLK) that may control WNT signaling and adhesion/cell migration in mesenchymal-like NB cells, respectively (**Fig. 3E** and **S3A**, and **Table S2**). On the other hand, CDK7 and BRD2 emerged as additional kinases that promote the noradrenergic neuronal-like NB cell phenotype at the transcriptional level (**Fig. 3E** and **S3A**, and **Table S2**).

Despite 2^nd^ gen kiCCA identifying more than 6-times as many kinase interactions as our 1^st^ gen approach, one shortcoming of the approach is that we use cell and tissue lysates for our analyses. We speculate that we lose most kinase PPI observations because cellular contents are diluted by about 100-fold upon cell lysis, which will lead to the dissociation of low-affinity PPIs; this favors the identification of stable kinase complexes and disfavors the identification of kinase-substrate interactions. The next generations of kiCCA will overcome these shortcomings by capturing kinase complexes in intact cells and tissues.

## ACKNOWLEDGMENTS

This work was supported by grants from the National Institutes of Health issued under the award numbers R01GM129090 (S-E.O.), R03TR003308 (M.G.), 1R35GM150766 (M.G.), and P30CA042014 (M.G.). The content is solely the responsibility of the authors and does not necessarily represent the official views of the National Institutes of Health.

## AUTHOR CONTRIBUTIONS

Conceptualization, M.G.; Methodology, M.G.; Investigation, M.G., K.W., T.A.R., A.M.C., P.J., G.E.M., K.M., T.S.; Formal Analysis, M.G., K.O.; Writing – Original Draft, M.G.; Writing – Review and Editing, M.G., S-E.O., K.W., T.A.R, A.M.C; Funding Acquisition, S-E.O. and M.G.

## RESOURCE AVAILABILITY

### Lead Contact

Martin Golkowski, Department of Pharmacology & Toxicology, University of Utah, Salt Lake City, UT 84112, USA

### Materials Availability

As lead contact, Martin Golkowski is responsible for all reagent and resource requests. Please contact Martin Golkowski at with requests and inquiries.

### Data and Code Availability

Bruker MS output files, a detailed list of instrument settings, and DIA-NN output files generated by this study have been uploaded to the MassIVE repository of the University of San Diego under the acquisition number MSV000096379. This study did not generate new code.

## EXPERIMENTAL MODEL AND SUBJECT DETAILS

### Cell lines and tissue culture conditions

The neuroblastoma cell lines SK-N-SH and SH-SY5Y were purchased from the American Type Culture Collection (ATCC). All cells were grown at 37°C under 5% CO_2_, 95% ambient atmosphere. Fifteen cryo-frozen cell stocks were generated from the original vial from the cell bank (passage 3). Experiments were performed with cells at <10 passages from the original vial. All cell media used were those recommended by the ATCC, supplemented with 100x penicillin-streptomycin-glutamine (Thermo Fisher Scientific) and 10% fetal bovine serum (FBS, Corning). Cells were harvested when reaching 90% confluency.

### Trans well migration assay

For the trans well migration assay, 1*10E6 SH-SY5Y cells or SK-N-SH cells were seeded onto 6-well trans well inserts (Corning, 24-mm inserts, 8 µm pore size) in 1.35 mL of serum-free medium (see ‘*Cell lines and tissue culture conditions*’ above). Then either DMSO vehicle or drug in DMSO were added in 150 µL of serum free medium (10X stock) to reach a final DMSO concentration of 0.1% (v/v). Then inserts were placed in a 6-well plate well containing either 2.6 mL serum-free medium (negative control) or 2.6 mL complete growth medium containing 10% FBS (attractant). Cells were allowed to migrate for 24 h in a cell culture incubator at 37°C under 5% CO_2_, 95% ambient atmosphere. Then, inserts were gently washed with phosphate-buffered saline (PBS) twice and cells on top of the insert removed using a moist Q-tip. Wells were then placed in a new 6-well plate, where each well contained 1 mL of a 0.1% crystal violet solution in 20% aqueous methanol solution.

Cells were stained with crystal violet for 10 min at RT on a rocker and excess crystal violet solution was removed by rinsing the inserts twice with PBS. The inserts with the cells were then dried overnight at room temperature (RT). Then, using 70% ethanol and Kim wipes, any remaining crystal violet was removed from the top of the insert. Crystal violet-stained cells were then eluted into a new 6-well plate using 1 mL 33% aqueous acetic acid (v/v) on a rocker to allow mixing to homogeneity. 100 µL aliquots from each well were then transferred to standard clear flat bottom 96-well plates for quantification of the crystal violet concentration using a photo spectrometric plate reader at 590 nm wavelength (SkyHigh Plate Reader, Thermo Fisher Scientific).

The read-out was compared to a calibration curve obtained from a 96-well plate containing a defined number of cells. Drugs and concentrations used were the TBK1 inhibitor GSK8612 (2 µM final, Mechem Express, MCE), the CK2 inhibitor SGC-CK2-1 (2 µM final, MCE), and dasatinib (100 nM final, MCE).

### Inhibitor treatments for global proteome profiling

For global proteome profiling, 0.3x10^6^ SK-N-SH or SH-SY5Y cells per well were seeded on 6-well plates and allowed to adhere for 24 h. Then, the TBK1 inhibitor GSK8612 (2 µM final, MCE), the CK2 inhibitor SGC-CK2- 1 (2 µM final, MCE), or DMSO (vehicle control) were added. The final concentration of DMSO in all wells was 0.1% (v/v). Cells continued to grow for 96 h and then were harvested for global proteome analysis. Briefly, the growth medium was aspirated and the cells rinsed twice with ice-cold PBS. Cells were lysed in 8 M urea containing 100 mM Tris (pH =8.5), 5 mM tris(2-carboxyethyl)phosphine hydrochloride (TCEP*HCl) and 10 mM chloroacetamide (CAM). Lysates were harvested using a cell scraper (Sarstedt) and transferred to 1.7 mL microtubes and incubated on a thermal shaker for 30 min at 37°C and 1,400 rpm. For further sample processing, see ‘*Peptide preparation for global proteomics below’*.

### Kinase affinity enrichment, KI competition, and on-bead digestion of proteins

Kinase affinity enrichment, KI competition, and on-bead digestion of proteins was performed as previously described.^23^ In addition to the 21 KIPs, the following kinase inhibitors were used as the competitors for soluble competition experiments at the given final concentrations: lorlatinib (1 µM final, MCE), entrectinib (1 µM final, MCE), crizotinib (1 µM final, MCE), and ceritinib (1 µM final, MCE). Final DMSO (vehicle) concentration in all pulldowns was 0.1% (v/v).

### Co-immunoprecipitation/MS (Co-IP/MS) analyses of AP2B1 and AAK1 interactions

Co-IP/MS analyses were performed as previously described.^23^ Antibodies used were AAK1 (E8M3P) Rabbit mAb (Cell Signaling Technology, CST, #61527) and beta 2 Adaptin (AP2B1) pAb (Novus Biologicals, # NBP3- 29580). The resulting peptide samples were analyzed by nLC/MS on the Thermo Orbitrap Fusion Lumos system, and MS .raw files computed exactly as described previously.^23^

### Global proteome profiling

For global proteome analyses by nLC-MS, aliquots of 100 µg of protein in 8 M urea lysis buffer were pipetted into a new microtube and diluted with four times the volume ice-cold acetone. 80% aq. trichloroacetic acid (TCA) was added to a final concentration of 4% (v/v), samples were vortexed briefly at max speed and kept at - 20°C overnight. Precipitated protein was pelleted at 2,000 rcf at 4°C for 10 min and the supernatant aspirated. The pellet was cleaned by adding 1 mL of ice-cold acetone, dispersing the pellet in a sonicator bath, pelleting the protein at 2,000 rcf at 4°C for 10 min, and aspirating the supernatant; this step was repeated once more.

The pellets were dried until translucent (2-3 min), 100 µL of 8 M urea buffer containing 100 mM Tris (pH = 8.5) was added and the sample agitated on a thermal shaker at 1,400 rpm at 37°C until the pellet was dissolved (15-30 min). Then, the samples were diluted 2-fold with 100 mM triethylammonium bicarbonate (TEAB), the pH was adjusted to 8-9 with 1 M aq. NaOH, if needed, and 1 µg of Lys-C (Wako-Fujifilm) was added (1:100 ratio digestive enzyme to protein). Then, the mixtures were agitated on a thermal shaker at 1,400 rpm at 37°C for 2 h, diluted another 2-fold with 100 mM TEAB, and 1 µg of MS grade trypsin was added (Pierce). The pH was tested and readjusted if needed. The mixtures were agitated on a thermomixer at 1,400 rpm at 37°C overnight, acidified with formic acid (FA, 1% final), and cleared by centrifugation for 10 min at 14,000 rcf and RT. An aliquot of the supernatant equaling 5 µg peptide was desalted on C18 StageTips to be used for global proteome analysis by nLC-MS.^80^

### nLC/MS analyses of peptide samples on the Bruker timsTOF Pro 2 system

Peptide samples of 200 ng were analyzed on a timsTOF Pro 2 – nanoElute 2 nLC-MS system (Bruker). Peptides were separated using 15 cm long, 150 µm inner diameter PepSep columns packed with 1.5 µm diameter C_18_ beads in 45 min long LC gradients (5-30% B) at flow rates of 0.5 µL/min. LC solvents were A (0.1% formic acid in LC-MS-grade water) and B (0.1% formic acid in LC-MS-grade acetonitrile). MS data were acquired in data-independent acquisition mode using the diaPASEF method.^40^ General MS instrument parameters were the following: polarity: positive; mass range: 100 m/z to 1700 m/z; 1/K_0_ start: 0.60 Vs/cm²; 1/K_0_ end: 1.40 Vs/cm², CaptiveSpray; capillary: 1600 V. The complete list of instrument setting can be found in the MassIVE repository of the University of San Diego under the acquisition number MSV000096379.

### Computation of Bruker MS raw files and output data processing

Bruker MS raw files were computed using DIA-NN v.1.8.1 in library-free mode.^52^ The Homo sapiens FASTA downloaded from UniProt on 07.14.2023 (UP000005640) was used in the DIA-NN search. Briefly, DIA-NN search parameters were the following: reannotate: enabled; FASTA digest for library-free search/library generation: enabled; deep learning-based spectra, RTs, and IMs prediction: enabled; protease: Trypsin/P; missed cleavages: 1; number of variable modifications: 0; N-term M excision: enabled; C carbamidomethylation: enabled; peptide length range: 7 - 30; precursor charge range: 1 - 4; precursor m/z range: 300 - 1800; fragment ion m/z range: 200 – 1800; generate spectral library: enabled; quantities matrices: enabled; precursor FDR (%): 1.0; mass accuracy: 0.0; MS1 accuracy: 0.0; scan window: 0; use isotopologues: enabled; MBR: enabled; no shared spectra: enabled; protein inference: Genes; neural network classifier: Single pass mode; quantification strategy: Robust LC (high precision); cross-run normalization: global; smart profiling; speed and RAM usage: optimal results. DIA-NN raw output files were loaded into Perseus v2.0.10.0, log2 transformed, median-normalized, and missing values were imputed (width: 0.2; downshift 1.8).^81^ Perseus was also used for differential expression analysis.

### 2^nd^ Generation competition and competition correlation analysis (CCA)

For each cell line and condition tested, 21 KIP competition experiments and one DMSO control experiment were performed in biological duplicate, resulting in 44 kinobead pulldown and LC-MS experiments per condition/cell line. For input kinases into CCA, the following two sample t-test cut-off cutoff criteria were applied: p ≤ 0.05, log2 MS intensity FC ≥ 0.5 for the SH-N-SH cell line and p ≤ 0.01, log2 MS intensity FC ≥ 0.5 for the SH-SY5Y cell line. All non-kinase proteins that were quantified were used for CCA. We correlated MS intensity values of all selected kinases and all non-kinase proteins using Pearson moment correlation (n = 44). We determined the kinases which showed the 1^st^ through 10^th^ highest Pearson r-value and estimated %TDR values for kinase PPI predictions based on the percentage of previously reported kinase PPIs in the BioGRID interactome database^53^ in each Pearson’s r-value interval of 0.05 units (%FDR = (N_PPI,reported_ / N_PPI,unreported_) * 100)). For mapping of previously reported PPIs we used the interactions derived from the ‘BIOGRID-MV-Physical-4.4.223.mitab’ file downloaded on July 21, 2024. Predicted kinase-protein interactions showing a %TDR > 10 were then reported as a predicted PPIs with the %TDR value associated with each PPI (**Table S2**).

### Differential expression analysis (DEA)

To identify differentially expressed proteomic features between cell lines and treatment conditions, we applied either a two-sample Student’s t-test or a one sample Student’s t-test, applying Benjamini-Hochberg (BH) correction for multiple hypothesis testing (FDR ≤ 0.05, discovery mode in kinobead AP-MS experiments and to analyze 2^nd^ gen kiCCA data). Alternatively, we applied a simple p ≤ 0.05 (validation mode in kinobead and Co-IP/MS data).^23^

### Plotting STRING interaction networks

PPI network models were plotted using the STRING web application version 12.0 with the following settings: Edges were scaled with confidence, and only ‘physical subnetwork’ interactions were considered, i.e., only considering text mining, experiments, and databases.^82^

### Gene set enrichment analysis (GSEA)

**(A)** For gene set enrichment analysis (GSEA), we used the ssGSEA2.0 script in R together with the Gene Ontology: Biological Process (GOBP) gene set of the MSigDB database (‘c5.bp.v7.0.symbols’) according to the published protocol with the following minor modifications:^83^ to rank gene names, we calculated a compound score using the two sample Student’s t-test log2 MS intensity difference multiplied by the -log10 p-value. The parameters used for GSEA were: sample.norm.type = “none”, weight = 1, statistic = “area.under.RES”, output.score.type = “NES”, nperm = 1e3, min.overlap = 10, correl.type = “z.score”, par = T, spare.cores = 1, export.signat.gct = T, extended.output = T.

## SUPPLEMENTARY FIGURES

**Figure S1.**
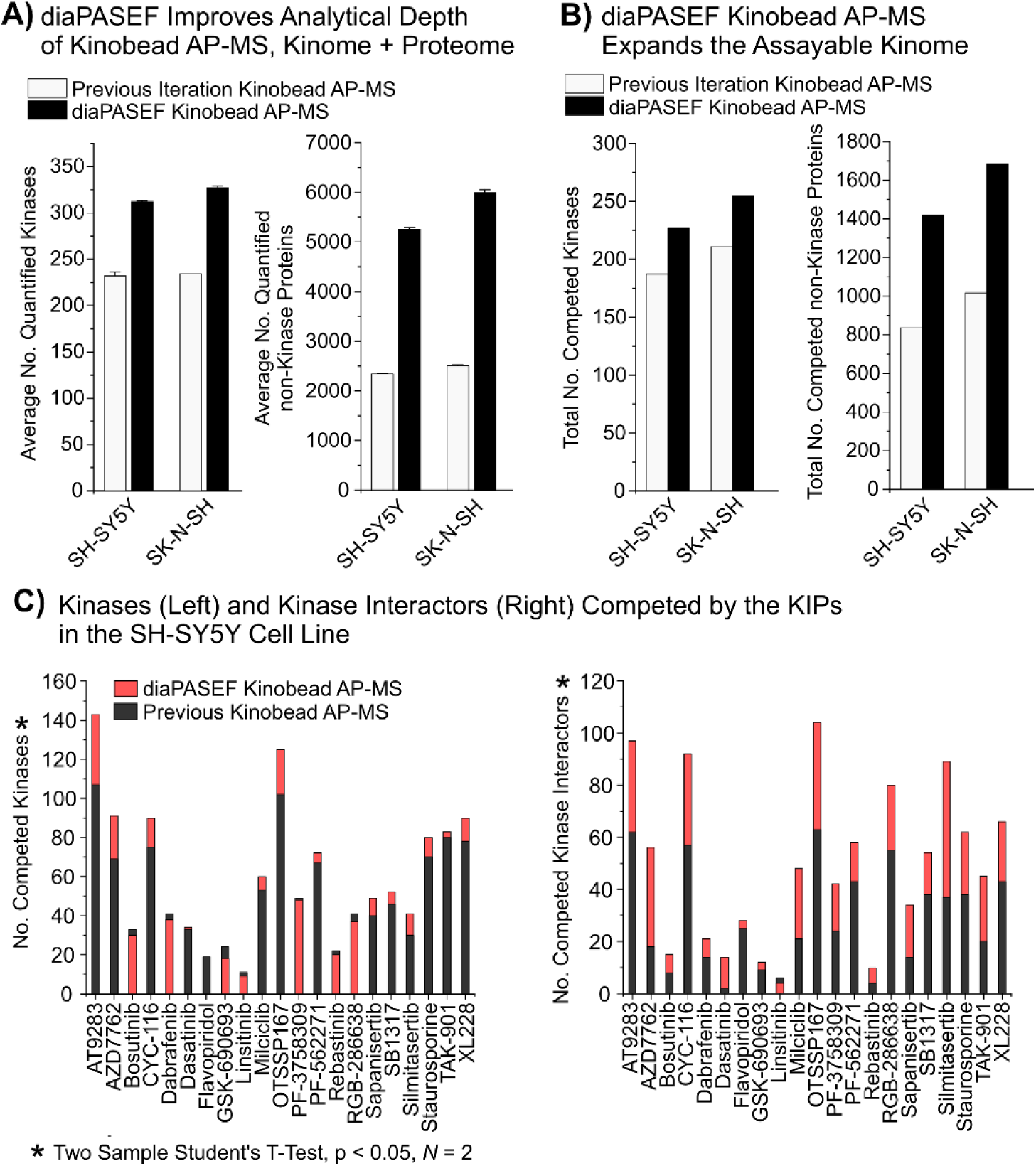
(A) Number of quantified protein kinases and non-kinase proteins in the SH-SY5Y and SK-N-SH cell lines, comparing our previous iteration of kinobead AP-MS with diaPASEF-powered kinobead AP-MS. Refers to Fig. 1B, and **Table S1**. **(B)** Number of quantified protein kinases and non-kinase proteins that could be competed with the 21 KIPs in the SH- SY5Y and SK-N-SH cell lines, comparing our previous iteration of kinobead AP-MS with diaPASEF kinobead AP-MS. Statistics: two-sample Student’s t-test p-value ≤ 0.05, log2 FC ≥ 0, *N* = 2. Refers for Fig. 1C and **Table S1**. **(C)** Number of kinases (left panel) and reported non-kinase interaction partners of these kinases (right panel) that were significantly competed against the kinobeads in the SH-SY5Y cell line with our 21 KIPs in our diaPASEF kinobead AP-MS experiments with soluble competition, compared to our previous experiments (two-sample Student’s t-test p-value < 0.05, log2 FC > 0, *N* = 2). Refers to Fig. 1C and **Table S1**.

**Figure S2.**
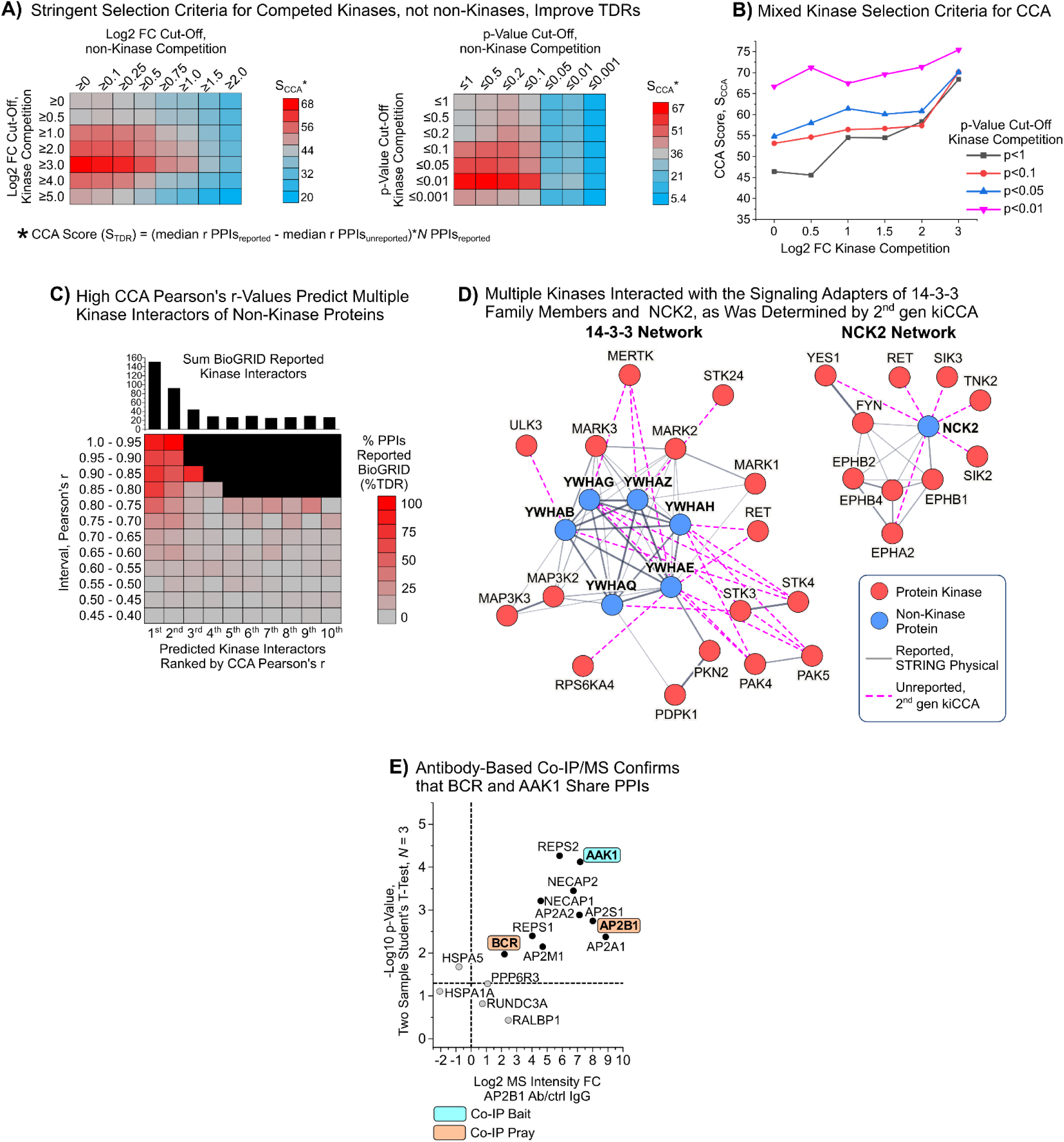
(A) Parameter scanning experiment determining the CCA score (SCCA) for kinase PPIs predicted by our 2^nd^ gen CCA in the SH-SY5Y cell line. Increasingly stringent Student’s t-test log2 FC and p-value cut-off criteria were applied and only kinases or non-kinases fulfilling these criteria used for CCA. Refers to Fig. 2A. (B) Plotting the SCCA for varying log2 FC cut-offs and p-Value cut-offs against one another identifies optimal mixed cut-off criteria for kinases input into our CCA of the SH-SY5Y cell line. Refers to Fig. 2B. (C) Heatmap showing the %TDR of our CCA for kinase interactions by 0.05-unit intervals of the CCA Pearson’s r-value for the 1^st^ through 10^th^ predicted kinase interactor of proteins. 2^nd^ gen CCA data from SH-SY5Y cells. Refers to Fig. 2D. (D) Physical STRING interaction network (v12.0) of multiple kinases predicted to interact with the signaling scaffolds NCK2 and 14-3-3 family proteins. Only interactions with a 2^nd^ gen kiCCA %TDR > 10 were used to construct the networks. (E) Volcano plot showing the results of a co-IP/MS experiment using a specific antibody that binds AAK1. Only proteins that are also members of the STRING interaction network in Fig. 2E are shown. Statistics: two sample Student’s t-test p < 0.05. Refers to Fig. 2E and **Table S2**.

**Figure S3.**
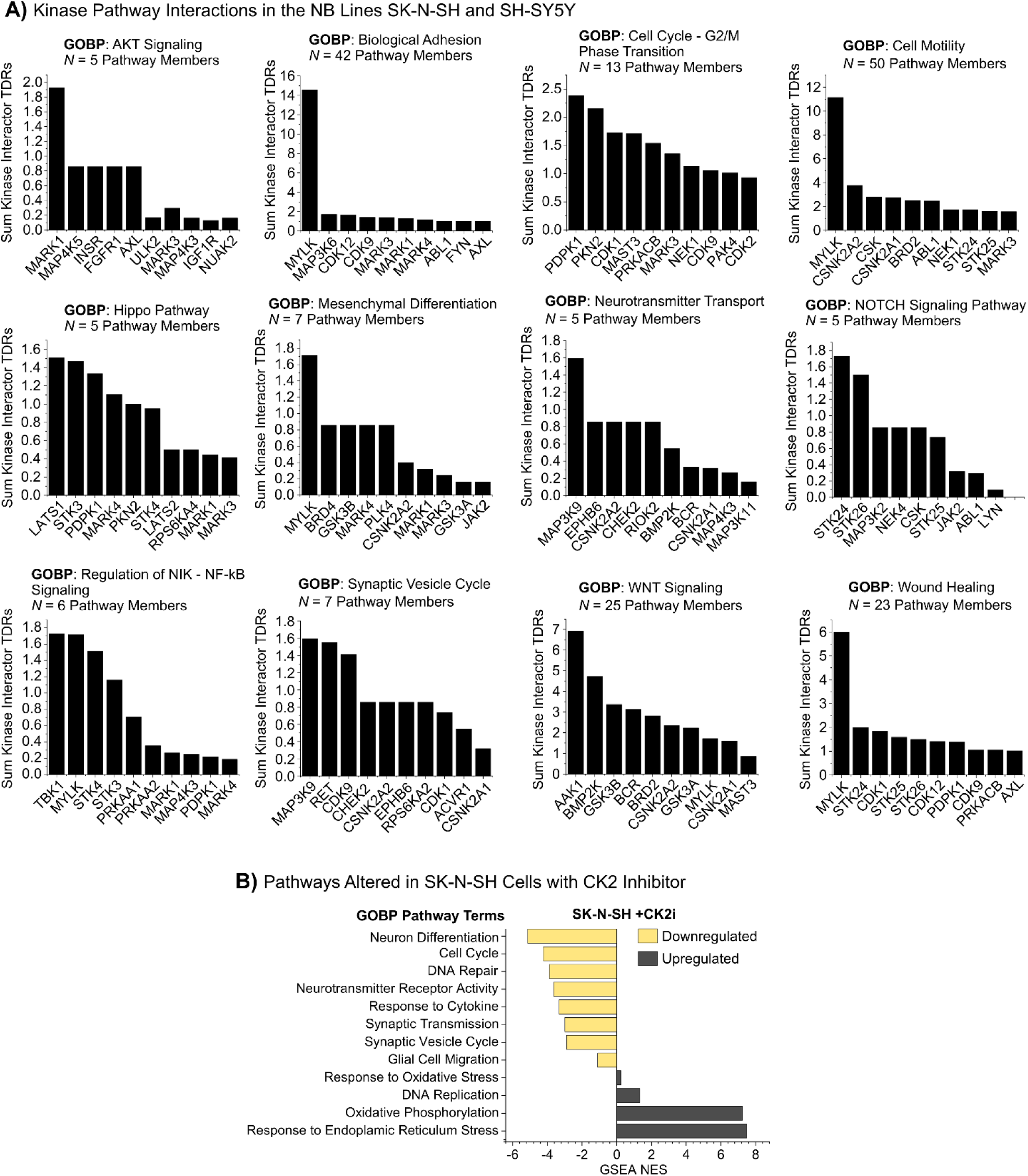
(A) Association of kinases with signaling pathways through their interaction partners. Kinases were ranked by the sum %TDRs for PPIs with their interaction partners. Interaction partners were deemed members of a signaling pathway if their gene name was contained in the GOBP gene sets shown in Fig. 3D. Refers to Fig. 3D and **Table S2**. Results from GSEA with GOBP terms of global proteomics expression data, comparing SK-N-SH cells treated with either the CK2 inhibitor SGC-CK2-1 or DMSO (vehicle) for 4 days. Only pathway terms that showed an FDR < 0.05 are shown. Relates to and **Table S2**.

## REFERENCES

1. Ideker T, Sharan R. Protein networks in disease. Genome Res. 2008;18(4):644–52. doi: 10.1101/gr.071852.107. PubMed PMID: 18381899; PMCID: PMC3863981.

2. Nussinov R. The spatial structure of cell signaling systems. Phys Biol. 2013;10(4):045004. doi: 10.1088/1478-3975/10/4/045004. PubMed PMID: 23913102; PMCID: PMC3870181.

3. Richards AL, Eckhardt M, Krogan NJ. Mass spectrometry-based protein-protein interaction networks for the study of human diseases. Mol Syst Biol. 2021;17(1):e8792. doi: 10.15252/msb.20188792. PubMed PMID: 33434350; PMCID: PMC7803364.

4. Cheng F, Zhao J, Wang Y, Lu W, Liu Z, Zhou Y, Martin WR, Wang R, Huang J, Hao T, Yue H, Ma J, Hou Y, Castrillon JA, Fang J, Lathia JD, Keri RA, Lightstone FC, Antman EM, Rabadan R, Hill DE, Eng C, Vidal M, Loscalzo J. Comprehensive characterization of protein-protein interactions perturbed by disease mutations. Nat Genet. 2021;53(3):342–53. doi: 10.1038/s41588-020-00774-y. PubMed PMID: 33558758; PMCID: PMC8237108.

5. Huttlin EL, Bruckner RJ, Paulo JA, Cannon JR, Ting L, Baltier K, Colby G, Gebreab F, Gygi MP, Parzen H, Szpyt J, Tam S, Zarraga G, Pontano-Vaites L, Swarup S, White AE, Schweppe DK, Rad R, Erickson BK, Obar RA, Guruharsha KG, Li K, Artavanis-Tsakonas S, Gygi SP, Harper JW. Architecture of the human interactome defines protein communities and disease networks. Nature. 2017;545(7655):505–9. doi: 10.1038/nature22366. PubMed PMID: 28514442; PMCID: PMC5531611.

6. Taylor IW, Wrana JL. Protein interaction networks in medicine and disease. Proteomics. 2012;12(10):1706–16. doi: 10.1002/pmic.201100594. PubMed PMID: 22593007.

7. Duan G, Walther D. The roles of post-translational modifications in the context of protein interaction networks. PLoS Comput Biol. 2015;11(2):e1004049. doi: 10.1371/journal.pcbi.1004049. PubMed PMID: 25692714; PMCID: PMC4333291.

8. Vidal M, Cusick ME, Barabasi AL. Interactome networks and human disease. Cell. 2011;144(6):986-98. doi: 10.1016/j.cell.2011.02.016. PubMed PMID: 21414488; PMCID: PMC3102045.

9. Manning G, Whyte DB, Martinez R, Hunter T, Sudarsanam S. The protein kinase complement of the human genome. Science. 2002;298(5600):1912–34. doi: 10.1126/science.1075762. PubMed PMID: 12471243.

10. Fleuren ED, Zhang L, Wu J, Daly RJ. The kinome ’at large’ in cancer. Nat Rev Cancer. 2016;16(2):83–98. doi: 10.1038/nrc.2015.18. PubMed PMID: 26822576.

11. Ferguson FM, Gray NS. Kinase inhibitors: the road ahead. Nat Rev Drug Discov. 2018;17(5):353–77. doi: 10.1038/nrd.2018.21. PubMed PMID: 29545548.

12. Cohen P, Cross D, Janne PA. Kinase drug discovery 20 years after imatinib: progress and future directions. Nat Rev Drug Discov. 2021;20(7):551–69. doi: 10.1038/s41573-021-00195-4. PubMed PMID: 34002056; PMCID: PMC8127496 member of the Scientific Advisory Boards of Mission Therapeutics, Ubiquigent and Biocatalyst International. D.C. is an employee and shareholder of AstraZeneca. P.A.J. has received consulting fees from AstraZeneca, Boehringer-Ingelheim, Pfizer, Roche/Genentech, Takeda Oncology, ACEA Biosciences, Eli Lilly and Company, Araxes Pharma, Ignyta, Mirati Therapeutics, Novartis, LOXO Oncology, Daiichi Sankyo, Sanofi Oncology, Voronoi, SFJ Pharmaceuticals, Biocartis, Novartis Oncology, Nuvalent, Esai, Bayer, Transcenta and Silicon Therapeutics; receives post-marketing royalties from DFCI-owned intellectual property on EGFR mutations licensed to Lab Corp; has sponsored research agreements with AstraZeneca, Daichi- Sankyo, PUMA, Boehringer-Ingelheim, Eli Lilly and Company, Revolution Medicines, and Astellas Pharmaceuticals; and has stock ownership in Gatekeeper Pharmaceuticals.

13. Attwood MM, Fabbro D, Sokolov AV, Knapp S, Schioth HB. Trends in kinase drug discovery: targets, indications and inhibitor design. Nat Rev Drug Discov. 2021;20(11):839–61. doi: 10.1038/s41573-021-00252-y. PubMed PMID: 34354255.

14. Bhullar KS, Lagaron NO, McGowan EM, Parmar I, Jha A, Hubbard BP, Rupasinghe HPV. Kinase-targeted cancer therapies: progress, challenges and future directions. Mol Cancer. 2018;17(1):48. doi: 10.1186/s12943-018-0804-2. PubMed PMID: 29455673; PMCID: PMC5817855.

15. Zarrin AA, Bao K, Lupardus P, Vucic D. Kinase inhibition in autoimmunity and inflammation. Nat Rev Drug Discov. 2021;20(1):39–63. doi: 10.1038/s41573-020-0082-8. PubMed PMID: 33077936; PMCID: PMC7569567 K.B. is an employee of Genentech. P.L. is an employee of Synthekine and holds stock in Synthekine and the Roche Group. D.V. is an employee of Genentech and holds stock and options in the Roche Group.

16. Castelo-Soccio L, Kim H, Gadina M, Schwartzberg PL, Laurence A, O’Shea JJ. Protein kinases: drug targets for immunological disorders. Nat Rev Immunol. 2023;23(12):787-806. doi: 10.1038/s41577-023-00877-7. PubMed PMID: 37188939; PMCID: PMC10184645 immune modulators’ US7070972B1. J.J.O’S. receives income from the US Government for royalties>. All other authors have no competing interests.

17. Chico LK, Van Eldik LJ, Watterson DM. Targeting protein kinases in central nervous system disorders. Nat Rev Drug Discov. 2009;8(11):892–909. doi: 10.1038/nrd2999. PubMed PMID: 19876042; PMCID: PMC2825114.

18. Garcia-Carceles J, Caballero E, Gil C, Martinez A. Kinase Inhibitors as Underexplored Antiviral Agents. J Med Chem. 2022;65(2):935–54. doi: 10.1021/acs.jmedchem.1c00302. PubMed PMID: 33970631; PMCID: PMC8802305.

19. Arendse LB, Wyllie S, Chibale K, Gilbert IH. Plasmodium Kinases as Potential Drug Targets for Malaria: Challenges and Opportunities. ACS Infect Dis. 2021;7(3):518–34. doi: 10.1021/acsinfecdis.0c00724. PubMed PMID: 33590753; PMCID: PMC7961706.

20. Lemmon MA, Schlessinger J. Cell signaling by receptor tyrosine kinases. Cell. 2010;141(7):1117–34. doi: 10.1016/j.cell.2010.06.011. PubMed PMID: 20602996; PMCID: PMC2914105.

21. Pawson T, Scott JD. Signaling through scaffold, anchoring, and adaptor proteins. Science. 1997;278(5346):2075–80. doi: 10.1126/science.278.5346.2075. PubMed PMID: 9405336.

22. Barbour H, Nkwe NS, Estavoyer B, Messmer C, Gushul-Leclaire M, Villot R, Uriarte M, Boulay K, Hlayhel S, Farhat B, Milot E, Mallette FA, Daou S, Affar EB. An inventory of crosstalk between ubiquitination and other post-translational modifications in orchestrating cellular processes. iScience. 2023;26(5):106276. doi: 10.1016/j.isci.2023.106276. PubMed PMID: 37168555; PMCID: PMC10165199.

23. Golkowski M, Lius A, Sapre T, Lau HT, Moreno T, Maly DJ, Ong SE. Multiplexed kinase interactome profiling quantifies cellular network activity and plasticity. Mol Cell. 2023;83(5):803–18 e8. doi: 10.1016/j.molcel.2023.01.015. PubMed PMID: 36736316.

24. Golkowski M, Lau HT, Chan M, Kenerson H, Vidadala VN, Shoemaker A, Maly DJ, Yeung RS, Gujral TS, Ong SE. Pharmacoproteomics Identifies Kinase Pathways that Drive the Epithelial-Mesenchymal Transition and Drug Resistance in Hepatocellular Carcinoma. Cell Syst. 2020;11(2):196–207 e7. doi: 10.1016/j.cels.2020.07.006. PubMed PMID: 32755597; PMCID: PMC7484106.

25. Stynen B, Tournu H, Tavernier J, Van Dijck P. Diversity in genetic in vivo methods for protein-protein interaction studies: from the yeast two-hybrid system to the mammalian split-luciferase system. Microbiol Mol Biol Rev. 2012;76(2):331–82. doi: 10.1128/MMBR.05021-11. PubMed PMID: 22688816; PMCID: PMC3372256.

26. Buljan M, Ciuffa R, van Drogen A, Vichalkovski A, Mehnert M, Rosenberger G, Lee S, Varjosalo M, Pernas LE, Spegg V, Snijder B, Aebersold R, Gstaiger M. Kinase Interaction Network Expands Functional and Disease Roles of Human Kinases. Mol Cell. 2020;79(3):504–20 e9. doi: 10.1016/j.molcel.2020.07.001. PubMed PMID: 32707033; PMCID: PMC7427327.

27. Huttlin EL, Bruckner RJ, Navarrete-Perea J, Cannon JR, Baltier K, Gebreab F, Gygi MP, Thornock A, Zarraga G, Tam S, Szpyt J, Gassaway BM, Panov A, Parzen H, Fu S, Golbazi A, Maenpaa E, Stricker K, Guha Thakurta S, Zhang T, Rad R, Pan J, Nusinow DP, Paulo JA, Schweppe DK, Vaites LP, Harper JW, Gygi SP. Dual proteome-scale networks reveal cell-specific remodeling of the human interactome. Cell. 2021;184(11):3022–40 e28. doi: 10.1016/j.cell.2021.04.011. PubMed PMID: 33961781; PMCID: PMC8165030.

28. Kristensen AR, Gsponer J, Foster LJ. A high-throughput approach for measuring temporal changes in the interactome. Nat Methods. 2012;9(9):907–9. doi: 10.1038/nmeth.2131. PubMed PMID: 22863883; PMCID: PMC3954081.

29. O’Reilly FJ, Rappsilber J. Cross-linking mass spectrometry: methods and applications in structural, molecular and systems biology. Nat Struct Mol Biol. 2018;25(11):1000–8. doi: 10.1038/s41594-018-0147-0. PubMed PMID: 30374081.

30. Roux KJ, Kim DI, Raida M, Burke B. A promiscuous biotin ligase fusion protein identifies proximal and interacting proteins in mammalian cells. J Cell Biol. 2012;196(6):801–10. doi: 10.1083/jcb.201112098. PubMed PMID: 22412018; PMCID: PMC3308701.

31. Goos H, Kinnunen M, Salokas K, Tan Z, Liu X, Yadav L, Zhang Q, Wei GH, Varjosalo M. Human transcription factor protein interaction networks. Nat Commun. 2022;13(1):766. doi: 10.1038/s41467-022-28341-5. PubMed PMID: 35140242; PMCID: PMC8828895.

32. Rhee HW, Zou P, Udeshi ND, Martell JD, Mootha VK, Carr SA, Ting AY. Proteomic mapping of mitochondria in living cells via spatially restricted enzymatic tagging. Science. 2013;339(6125):1328–31. doi: 10.1126/science.1230593. PubMed PMID: 23371551; PMCID: PMC3916822.

33. Qin W, Cho KF, Cavanagh PE, Ting AY. Deciphering molecular interactions by proximity labeling. Nat Methods. 2021;18(2):133–43. doi: 10.1038/s41592-020-01010-5. PubMed PMID: 33432242.

34. Bantscheff M, Eberhard D, Abraham Y, Bastuck S, Boesche M, Hobson S, Mathieson T, Perrin J, Raida M, Rau C, Reader V, Sweetman G, Bauer A, Bouwmeester T, Hopf C, Kruse U, Neubauer G, Ramsden N, Rick J, Kuster B, Drewes G. Quantitative chemical proteomics reveals mechanisms of action of clinical ABL kinase inhibitors. Nat Biotechnol. 2007;25(9):1035–44. doi: 10.1038/nbt1328. PubMed PMID: 17721511.

35. Duncan JS, Whittle MC, Nakamura K, Abell AN, Midland AA, Zawistowski JS, Johnson NL, Granger DA, Jordan NV, Darr DB, Usary J, Kuan PF, Smalley DM, Major B, He X, Hoadley KA, Zhou B, Sharpless NE, Perou CM, Kim WY, Gomez SM, Chen X, Jin J, Frye SV, Earp HS, Graves LM, Johnson GL. Dynamic reprogramming of the kinome in response to targeted MEK inhibition in triple-negative breast cancer. Cell. 2012;149(2):307–21. doi: 10.1016/j.cell.2012.02.053. PubMed PMID: 22500798; PMCID: PMC3328787.

36. Klaeger S, Heinzlmeir S, Wilhelm M, Polzer H, Vick B, Koenig PA, Reinecke M, Ruprecht B, Petzoldt S, Meng C, Zecha J, Reiter K, Qiao H, Helm D, Koch H, Schoof M, Canevari G, Casale E, Depaolini SR, Feuchtinger A, Wu Z, Schmidt T, Rueckert L, Becker W, Huenges J, Garz AK, Gohlke BO, Zolg DP, Kayser G, Vooder T, Preissner R, Hahne H, Tonisson N, Kramer K, Gotze K, Bassermann F, Schlegl J, Ehrlich HC, Aiche S, Walch A, Greif PA, Schneider S, Felder ER, Ruland J, Medard G, Jeremias I, Spiekermann K, Kuster B. The target landscape of clinical kinase drugs. Science. 2017;358(6367). doi: 10.1126/science.aan4368. PubMed PMID: 29191878; PMCID: PMC6542668.

37. Medard G, Pachl F, Ruprecht B, Klaeger S, Heinzlmeir S, Helm D, Qiao H, Ku X, Wilhelm M, Kuehne T, Wu Z, Dittmann A, Hopf C, Kramer K, Kuster B. Optimized chemical proteomics assay for kinase inhibitor profiling. J Proteome Res. 2015;14(3):1574–86. doi: 10.1021/pr5012608. PubMed PMID: 25660469.

38. Golkowski M, Vidadala VN, Lau HT, Shoemaker A, Shimizu-Albergine M, Beavo J, Maly DJ, Ong SE. Kinobead/LC-MS Phosphokinome Profiling Enables Rapid Analyses of Kinase-Dependent Cell Signaling Networks. J Proteome Res. 2020;19(3):1235–47. doi: 10.1021/acs.jproteome.9b00742. PubMed PMID: 32037842; PMCID: PMC7537592.

39. Golkowski M, Vidadala RS, Lombard CK, Suh HW, Maly DJ, Ong SE. Kinobead and Single-Shot LC-MS Profiling Identifies Selective PKD Inhibitors. J Proteome Res. 2017;16(3):1216–27. doi: 10.1021/acs.jproteome.6b00817. PubMed PMID: 28102076; PMCID: PMC5663466.

40. Meier F, Brunner AD, Frank M, Ha A, Bludau I, Voytik E, Kaspar-Schoenefeld S, Lubeck M, Raether O, Bache N, Aebersold R, Collins BC, Rost HL, Mann M. diaPASEF: parallel accumulation-serial fragmentation combined with data-independent acquisition. Nat Methods. 2020;17(12):1229–36. doi: 10.1038/s41592-020-00998-0. PubMed PMID: 33257825.

41. Matthay KK, Maris JM, Schleiermacher G, Nakagawara A, Mackall CL, Diller L, Weiss WA. Neuroblastoma. Nat Rev Dis Primers. 2016;2:16078. doi: 10.1038/nrdp.2016.78. PubMed PMID: 27830764.

42. Thirant C, Peltier A, Durand S, Kramdi A, Louis-Brennetot C, Pierre-Eugene C, Gautier M, Costa A, Grelier A, Zaidi S, Gruel N, Jimenez I, Lapouble E, Pierron G, Sitbon D, Brisse HJ, Gauthier A, Freneaux P, Grossetete S, Baudrin LG, Raynal V, Baulande S, Bellini A, Bhalshankar J, Carcaboso AM, Geoerger B, Rohrer H, Surdez D, Boeva V, Schleiermacher G, Delattre O, Janoueix-Lerosey I. Reversible transitions between noradrenergic and mesenchymal tumor identities define cell plasticity in neuroblastoma. Nat Commun. 2023;14(1):2575. doi: 10.1038/s41467-023-38239-5. PubMed PMID: 37142597; PMCID: PMC10160107.

43. Gautier M, Thirant C, Delattre O, Janoueix-Lerosey I. Plasticity in Neuroblastoma Cell Identity Defines a Noradrenergic-to-Mesenchymal Transition (NMT). Cancers (Basel). 2021;13(12). doi: 10.3390/cancers13122904. PubMed PMID: 34200747; PMCID: PMC8230375.

44. van Groningen T, Koster J, Valentijn LJ, Zwijnenburg DA, Akogul N, Hasselt NE, Broekmans M, Haneveld F, Nowakowska NE, Bras J, van Noesel CJM, Jongejan A, van Kampen AH, Koster L, Baas F, van Dijk-Kerkhoven L, Huizer-Smit M, Lecca MC, Chan A, Lakeman A, Molenaar P, Volckmann R, Westerhout EM, Hamdi M, van Sluis PG, Ebus ME, Molenaar JJ, Tytgat GA, Westerman BA, van Nes J, Versteeg R. Neuroblastoma is composed of two super-enhancer-associated differentiation states. Nat Genet. 2017;49(8):1261–6. doi: 10.1038/ng.3899. PubMed PMID: 28650485.

45. Villalard B, Boltjes A, Reynaud F, Imbaud O, Thoinet K, Timmerman I, Croze S, Theoulle E, Atzeni G, Lachuer J, Molenaar JJ, Tytgat GAM, Delloye-Bourgeois C, Castellani V. Neuroblastoma plasticity during metastatic progression stems from the dynamics of an early sympathetic transcriptomic trajectory. Nat Commun. 2024;15(1):9570. Epub 20241106. doi: 10.1038/s41467-024-53776-3. PubMed PMID: 39500881; PMCID: PMC11538482.

46. Ben Amar D, Thoinet K, Villalard B, Imbaud O, Costechareyre C, Jarrosson L, Reynaud F, Novion Ducassou J, Coute Y, Brunet JF, Combaret V, Corradini N, Delloye-Bourgeois C, Castellani V. Environmental cues from neural crest derivatives act as metastatic triggers in an embryonic neuroblastoma model. Nat Commun. 2022;13(1):2549. Epub 20220510. doi: 10.1038/s41467-022-30237-3. PubMed PMID: 35538114; PMCID: PMC9091272.

47. Naiditch JA, Jie C, Lautz TB, Yu S, Clark S, Voronov D, Chu F, Madonna MB. Mesenchymal change and drug resistance in neuroblastoma. J Surg Res. 2015;193(1):279–88. doi: 10.1016/j.jss.2014.07.018. PubMed PMID: 25128389.

48. Westerhout EM, Hamdi M, Stroeken P, Nowakowska NE, Lakeman A, van Arkel J, Hasselt NE, Bleijlevens B, Akogul N, Haneveld F, Chan A, van Sluis P, Zwijnenburg D, Volckmann R, van Noesel CJM, Adameyko I, van Groningen T, Koster J, Valentijn LJ, van Nes J, Versteeg R. Mesenchymal-Type Neuroblastoma Cells Escape ALK Inhibitors. Cancer Res. 2022;82(3):484–96. doi: 10.1158/0008-5472.CAN-21-1621. PubMed PMID: 34853072.

49. Mabe NW, Huang M, Dalton GN, Alexe G, Schaefer DA, Geraghty AC, Robichaud AL, Conway AS, Khalid D, Mader MM, Belk JA, Ross KN, Sheffer M, Linde MH, Ly N, Yao W, Rotiroti MC, Smith BAH, Wernig M, Bertozzi CR, Monje M, Mitsiades CS, Majeti R, Satpathy AT, Stegmaier K, Majzner RG. Transition to a mesenchymal state in neuroblastoma confers resistance to anti-GD2 antibody via reduced expression of ST8SIA1. Nat Cancer. 2022;3(8):976–93. doi: 10.1038/s43018-022-00405-x. PubMed PMID: 35817829; PMCID: PMC10071839.

50. Sengupta S, Das S, Crespo AC, Cornel AM, Patel AG, Mahadevan NR, Campisi M, Ali AK, Sharma B, Rowe JH, Huang H, Debruyne DN, Cerda ED, Krajewska M, Dries R, Chen M, Zhang S, Soriano L, Cohen MA, Versteeg R, Jaenisch R, Spranger S, Romee R, Miller BC, Barbie DA, Nierkens S, Dyer MA, Lieberman J, George RE. Mesenchymal and adrenergic cell lineage states in neuroblastoma possess distinct immunogenic phenotypes. Nat Cancer. 2022;3(10):1228–46. doi: 10.1038/s43018-022-00427-5. PubMed PMID: 36138189; PMCID: PMC10171398.

51. Cox J, Hein MY, Luber CA, Paron I, Nagaraj N, Mann M. Accurate proteome-wide label-free quantification by delayed normalization and maximal peptide ratio extraction, termed MaxLFQ. Mol Cell Proteomics. 2014;13(9):2513–26. doi: 10.1074/mcp.M113.031591. PubMed PMID: 24942700; PMCID: PMC4159666.

52. Demichev V, Messner CB, Vernardis SI, Lilley KS, Ralser M. DIA-NN: neural networks and interference correction enable deep proteome coverage in high throughput. Nat Methods. 2020;17(1):41–4. doi: 10.1038/s41592-019-0638-x. PubMed PMID: 31768060; PMCID: PMC6949130.

53. . Oughtred R, Rust J, Chang C, Breitkreutz BJ, Stark C, Willems A, Boucher L, Leung G, Kolas N, Zhang F, Dolma S, Coulombe-Huntington J, Chatr-Aryamontri A, Dolinski K, Tyers M. The BioGRID database: A comprehensive biomedical resource of curated protein, genetic, and chemical interactions. Protein Sci. 2021;30(1):187–200. doi: 10.1002/pro.3978. PubMed PMID: 33070389; PMCID: PMC7737760.

54. Kielbowski K, Zychowska J, Becht R. Anaplastic lymphoma kinase inhibitors-a review of anticancer properties, clinical efficacy, and resistance mechanisms. Front Pharmacol. 2023;14:1285374. doi: 10.3389/fphar.2023.1285374. PubMed PMID: 37954850; PMCID: PMC10634320.

55. Zia V, Lengyel CG, Tajima CC, de Mello RA. Advancements of ALK inhibition of non-small cell lung cancer: a literature review. Transl Lung Cancer Res. 2023;12(7):1563–74. doi: 10.21037/tlcr-22-619. PubMed PMID: 37577315; PMCID: PMC10413028.

56. Goldsmith KC, Park JR, Kayser K, Malvar J, Chi YY, Groshen SG, Villablanca JG, Krytska K, Lai LM, Acharya PT, Goodarzian F, Pawel B, Shimada H, Ghazarian S, States L, Marshall L, Chesler L, Granger M, Desai AV, Mody R, Morgenstern DA, Shusterman S, Macy ME, Pinto N, Schleiermacher G, Vo K, Thurm HC, Chen J, Liyanage M, Peltz G, Matthay KK, Berko ER, Maris JM, Marachelian A, Mosse YP. Lorlatinib with or without chemotherapy in ALK-driven refractory/relapsed neuroblastoma: phase 1 trial results. Nat Med. 2023;29(5):1092–102. doi: 10.1038/s41591-023-02297-5. PubMed PMID: 37012551; PMCID: PMC10202811 investigator of the NANT phase 1 trial of lorlatinib and is a consultant for Pfizer. Y.P.M. has previously received research funding from Pfizer and Novartis. Y.P.M. has also served as a consultant for Lilly, Auron Therapeutics and Jumo Health. Y.P.M. serves as a member of the Data and Safety Monitoring Committee for the ASCO TAPUR study and receives honoraria for this role. H.T. is an employee of Pfizer and holds Pfizer stock. J.S. was an employee of Pfizer during the conduct and analysis of this work, currently owns Pfizer stock and is currently employed by Roche-Genentech. M.L. is a postdoctoral fellow with funding supported by Pfizer. G.P., at the time of study, was employed by Pfizer and held company stock. A.M. and NANT clinical research operations received funding from Pfizer in support of this study. K.C.G. has been an uncompensated consultant to Y-mAbs Therapeutics. L.M. has been a consultant or advisor for Bayer, Bristol Myers Squibb, Illumina and Tesaro and served as an EDMC member on studies run by Eisai and Merck. A.D. has stock ownership in Pfizer and Viatris and has acted as a paid consultant or advisor for Merck, Ology Medical Education, Y-mAbs Therapeutics and GlaxoSmithKline. D.A.M. has been a consultant or advisory board member to Y-mAbs Therapeutics, Clarity Pharmaceuticals, RayzeBio and Oncoheroes Biosciences. G.S. has received research funding from Bristol Myers Squibb, MSDavenir, Roche and Pfizer for research projects distinct from this trial. K.M. is a consultant with Y-mabs Therapeutics and RayzeBio. R.M. is on the Data and Safety Monitoring Committee for Y-Mabs Therapeutics and Jubilant Draximage. M.M. has stock in Johnson & Johnson, GE Healthcare and Varian Medical Systems. M.M. has been a consultant or advisory board memebr to Y-mAbs Therapeutics and has received research funding from Bayer, Ignyta, Roche, Lilly, Merck, Oncternal Therapeutics, AbbVie, Jubilant Draxlmage and Actuate Therapeutics. The remaining authors declare no competing interests.

57. Iyer R, Wehrmann L, Golden RL, Naraparaju K, Croucher JL, MacFarland SP, Guan P, Kolla V, Wei G, Cam N, Li G, Hornby Z, Brodeur GM. Entrectinib is a potent inhibitor of Trk-driven neuroblastomas in a xenograft mouse model. Cancer Lett. 2016;372(2):179–86. doi: 10.1016/j.canlet.2016.01.018. PubMed PMID: 26797418; PMCID: PMC4792275.

58. Marsilje TH, Pei W, Chen B, Lu W, Uno T, Jin Y, Jiang T, Kim S, Li N, Warmuth M, Sarkisova Y, Sun F, Steffy A, Pferdekamper AC, Li AG, Joseph SB, Kim Y, Liu B, Tuntland T, Cui X, Gray NS, Steensma R, Wan Y, Jiang J, Chopiuk G, Li J, Gordon WP, Richmond W, Johnson K, Chang J, Groessl T, He YQ, Phimister A, Aycinena A, Lee CC, Bursulaya B, Karanewsky DS, Seidel HM, Harris JL, Michellys PY. Synthesis, structure-activity relationships, and in vivo efficacy of the novel potent and selective anaplastic lymphoma kinase (ALK) inhibitor 5-chloro-N2-(2-isopropoxy-5-methyl-4-(piperidin-4- yl)phenyl)-N4-(2-(isopropylsulfonyl)phenyl)pyrimidine-2,4-diamine (LDK378) currently in phase 1 and phase 2 clinical trials. J Med Chem. 2013;56(14):5675–90. doi: 10.1021/jm400402q. PubMed PMID: 23742252.

59. Zhu C, Wei Y, Wei X. AXL receptor tyrosine kinase as a promising anti-cancer approach: functions, molecular mechanisms and clinical applications. Mol Cancer. 2019;18(1):153. doi: 10.1186/s12943-019-1090-3. PubMed PMID: 31684958; PMCID: PMC6827209.

60. Huelse JM, Fridlyand DM, Earp S, DeRyckere D, Graham DK. MERTK in cancer therapy: Targeting the receptor tyrosine kinase in tumor cells and the immune system. Pharmacol Ther. 2020;213:107577. doi: 10.1016/j.pharmthera.2020.107577. PubMed PMID: 32417270; PMCID: PMC9847360.

61. Johnson TW, Richardson PF, Bailey S, Brooun A, Burke BJ, Collins MR, Cui JJ, Deal JG, Deng YL, Dinh D, Engstrom LD, He M, Hoffman J, Hoffman RL, Huang Q, Kania RS, Kath JC, Lam H, Lam JL, Le PT, Lingardo L, Liu W, McTigue M, Palmer CL, Sach NW, Smeal T, Smith GL, Stewart AE, Timofeevski S, Zhu H, Zhu J, Zou HY, Edwards MP. Discovery of (10R)-7-amino-12-fluoro-2,10,16-trimethyl-15-oxo- 10,15,16,17-tetrahydro-2H-8,4-(metheno)pyrazolo[4,3-h][2,5,11]-benzoxadiazacyclotetradecine-3- carbonitrile (PF-06463922), a macrocyclic inhibitor of anaplastic lymphoma kinase (ALK) and c-ros oncogene 1 (ROS1) with preclinical brain exposure and broad-spectrum potency against ALK-resistant mutations. J Med Chem. 2014;57(11):4720-44. Epub 20140603. doi: 10.1021/jm500261q. PubMed PMID: 24819116.

62. Chan TY, Egbert CM, Maxson JE, Siddiqui A, Larsen LJ, Kohler K, Balasooriya ER, Pennington KL, Tsang TM, Frey M, Soderblom EJ, Geng H, Muschen M, Forostyan TV, Free S, Mercenne G, Banks CJ, Valdoz J, Whatcott CJ, Foulks JM, Bearss DJ, O’Hare T, Huang DCS, Christensen KA, Moody J, Warner SL, Tyner JW, Andersen JL. TNK1 is a ubiquitin-binding and 14-3-3-regulated kinase that can be targeted to block tumor growth. Nat Commun. 2021;12(1):5337. doi: 10.1038/s41467-021-25622-3. PubMed PMID: 34504101; PMCID: PMC8429728 Dainippon Pharma Oncology. Authors affiliated with Sumitomo Dainippon Pharma Oncology have a financial stake in the development of the TNK1 inhibitor. The remaining authors declare no competing interests.

63. Lee S, Rauch J, Kolch W. Targeting MAPK Signaling in Cancer: Mechanisms of Drug Resistance and Sensitivity. Int J Mol Sci. 2020;21(3). doi: 10.3390/ijms21031102. PubMed PMID: 32046099; PMCID: PMC7037308.

64. Sun C, Bernards R. Feedback and redundancy in receptor tyrosine kinase signaling: relevance to cancer therapies. Trends Biochem Sci. 2014;39(10):465–74. doi: 10.1016/j.tibs.2014.08.010. PubMed PMID: 25239057.

65. Cohen P. Signal integration at the level of protein kinases, protein phosphatases and their substrates. Trends Biochem Sci. 1992;17(10):408–13. doi: 10.1016/0968-0004(92)90010-7. PubMed PMID: 1333658.

66. Graves LM, Duncan JS, Whittle MC, Johnson GL. The dynamic nature of the kinome. Biochem J. 2013;450(1):1–8. doi: 10.1042/BJ20121456. PubMed PMID: 23343193; PMCID: PMC3808244.

67. Gupta PB, Pastushenko I, Skibinski A, Blanpain C, Kuperwasser C. Phenotypic Plasticity: Driver of Cancer Initiation, Progression, and Therapy Resistance. Cell Stem Cell. 2019;24(1):65–78. doi: 10.1016/j.stem.2018.11.011. PubMed PMID: 30554963; PMCID: PMC7297507.

68. Lamouille S, Xu J, Derynck R. Molecular mechanisms of epithelial-mesenchymal transition. Nat Rev Mol Cell Biol. 2014;15(3):178–96. doi: 10.1038/nrm3758. PubMed PMID: 24556840; PMCID: PMC4240281.

69. Gao Z, Lee P, Stafford JM, von Schimmelmann M, Schaefer A, Reinberg D. An AUTS2-Polycomb complex activates gene expression in the CNS. Nature. 2014;516(7531):349–54. doi: 10.1038/nature13921. PubMed PMID: 25519132; PMCID: PMC4323097.

70. Gao Z, Zhang J, Bonasio R, Strino F, Sawai A, Parisi F, Kluger Y, Reinberg D. PCGF homologs, CBX proteins, and RYBP define functionally distinct PRC1 family complexes. Mol Cell. 2012;45(3):344–56. doi: 10.1016/j.molcel.2012.01.002. PubMed PMID: 22325352; PMCID: PMC3293217.

71. Runde AP, Mack R, S JP, Zhang J. The role of TBK1 in cancer pathogenesis and anticancer immunity. J Exp Clin Cancer Res. 2022;41(1):135. doi: 10.1186/s13046-022-02352-y. PubMed PMID: 35395857; PMCID: PMC8994244.

72. Lavorgna A, De Filippi R, Formisano S, Leonardi A. TNF receptor-associated factor 1 is a positive regulator of the NF-kappaB alternative pathway. Mol Immunol. 2009;46(16):3278–82. doi: 10.1016/j.molimm.2009.07.029. PubMed PMID: 19698991.

73. Kato T, Jr., Gotoh Y, Hoffmann A, Ono Y. Negative regulation of constitutive NF-kappaB and JNK signaling by PKN1-mediated phosphorylation of TRAF1. Genes Cells. 2008;13(5):509–20. doi: 10.1111/j.1365-2443.2008.01182.x. PubMed PMID: 18429822.

74. Sahu SK, Garding A, Tiwari N, Thakurela S, Toedling J, Gebhard S, Ortega F, Schmarowski N, Berninger B, Nitsch R, Schmidt M, Tiwari VK. JNK-dependent gene regulatory circuitry governs mesenchymal fate. EMBO J. 2015;34(16):2162–81. doi: 10.15252/embj.201490693. PubMed PMID: 26157010; PMCID: PMC4557668.

75. Huber MA, Azoitei N, Baumann B, Grunert S, Sommer A, Pehamberger H, Kraut N, Beug H, Wirth T. NF-kappaB is essential for epithelial-mesenchymal transition and metastasis in a model of breast cancer progression. J Clin Invest. 2004;114(4):569–81. doi: 10.1172/JCI21358. PubMed PMID: 15314694; PMCID: PMC503772.

76. Kim CM, Choi JY, Bhat EA, Jeong JH, Son YJ, Kim S, Park HH. Crystal structure of TRAF1 TRAF domain and its implications in the TRAF1-mediated intracellular signaling pathway. Sci Rep. 2016;6:25526. doi: 10.1038/srep25526. PubMed PMID: 27151821; PMCID: PMC4858697.

77. Wells CI, Drewry DH, Pickett JE, Tjaden A, Kramer A, Muller S, Gyenis L, Menyhart D, Litchfield DW, Knapp S, Axtman AD. Development of a potent and selective chemical probe for the pleiotropic kinase CK2. Cell Chem Biol. 2021;28(4):546–58 e10. doi: 10.1016/j.chembiol.2020.12.013. PubMed PMID: 33484635; PMCID: PMC8864761.

78. Thomson DW, Poeckel D, Zinn N, Rau C, Strohmer K, Wagner AJ, Graves AP, Perrin J, Bantscheff M, Duempelfeld B, Kasparcova V, Ramanjulu JM, Pesiridis GS, Muelbaier M, Bergamini G. Discovery of GSK8612, a Highly Selective and Potent TBK1 Inhibitor. ACS Med Chem Lett. 2019;10(5):780-5. doi: 10.1021/acsmedchemlett.9b00027. PubMed PMID: 31097999; PMCID: PMC6512007 are employees or former employees of Cellzome GmbH and GlaxoSmithKline, and the company funded the work.

79. Thorburn A, Thamm DH, Gustafson DL. Autophagy and cancer therapy. Mol Pharmacol. 2014;85(6):830-8. Epub 20140226. doi: 10.1124/mol.114.091850. PubMed PMID: 24574520; PMCID: PMC4014668.

80. Rappsilber J, Mann M, Ishihama Y. Protocol for micro-purification, enrichment, pre-fractionation and storage of peptides for proteomics using StageTips. Nat Protoc. 2007;2(8):1896–906. doi: 10.1038/nprot.2007.261. PubMed PMID: 17703201.

81. Tyanova S, Temu T, Sinitcyn P, Carlson A, Hein MY, Geiger T, Mann M, Cox J. The Perseus computational platform for comprehensive analysis of (prote)omics data. Nat Methods. 2016;13(9):731–40. doi: 10.1038/nmeth.3901. PubMed PMID: 27348712.

82. Szklarczyk D, Kirsch R, Koutrouli M, Nastou K, Mehryary F, Hachilif R, Gable AL, Fang T, Doncheva NT, Pyysalo S, Bork P, Jensen LJ, von Mering C. The STRING database in 2023: protein-protein association networks and functional enrichment analyses for any sequenced genome of interest. Nucleic Acids Res. 2023;51(D1):D638–D46. doi: 10.1093/nar/gkac1000. PubMed PMID: 36370105; PMCID: PMC9825434.

83. Krug K, Mertins P, Zhang B, Hornbeck P, Raju R, Ahmad R, Szucs M, Mundt F, Forestier D, Jane-Valbuena J, Keshishian H, Gillette MA, Tamayo P, Mesirov JP, Jaffe JD, Carr SA, Mani DR. A Curated Resource for Phosphosite-specific Signature Analysis. Mol Cell Proteomics. 2019;18(3):576–93. doi: 10.1074/mcp.TIR118.000943. PubMed PMID: 30563849; PMCID: PMC6398202.

